# A *RIPOR2* in-frame deletion is a frequent and highly penetrant cause of adult-onset hearing loss

**DOI:** 10.1101/2019.12.20.884726

**Authors:** Suzanne E. de Bruijn, Jeroen J. Smits, Chang Liu, Cornelis P. Lanting, Andy J. Beynon, Joëlle Blankevoort, Jaap Oostrik, Wouter Koole, Erik de Vrieze, DOOFNL Consortium, Cor W.R.J. Cremers, Frans P. M. Cremers, Susanne Roosing, Helger G. Yntema, Henricus P.M. Kunst, Bo Zhao, Ronald J.E. Pennings, Hannie Kremer

## Abstract

Hearing loss is one of the most prevalent disabilities worldwide, and has a significant impact on quality of life. The adult-onset type of the condition is highly heritable but the genetic causes are largely unknown, which is in contrast to childhood-onset hearing loss. We identified an in-frame deletion of 12 nucleotides in *RIPOR2* as a highly penetrant cause of adult-onset progressive hearing loss that segregated as an autosomal dominant trait in 12 families from the Netherlands. Hearing loss associated with the deletion in 63 subjects displayed variable audiometric characteristics and an average age of onset of 30.6 years (SD 14.9 years, range 0-70 years). A functional effect of the *RIPOR2* variant was demonstrated by aberrant localization of the mutant RIPOR2 in the stereocilia of cochlear hair cells and failure to rescue morphological defects in RIPOR2-deficient hair cells, in contrast to the wildtype protein. Strikingly, the *RIPOR2* variant is present in 18 of 22,952 individuals not selected for hearing loss in the Southeast Netherlands. Collectively, these data demonstrate that an inherited form of adult-onset hearing loss is relatively common, with potentially thousands of individuals at risk in the Netherlands and beyond, which makes it an attractive target for developing a (genetic) therapy.

## INTRODUCTION

Hearing loss (HL) is one of the most prevalent disabilities worldwide (1) and genetic factors importantly contribute to this condition. So far, 118 genes have been associated with nonsyndromic forms of sensorineural HL and variants in these genes explain a significant part of subjects with an early onset of HL, i.e., congenital or in childhood (2-4). Our knowledge of the genetic architecture of adult-onset HL is limited despite a high heritability which is estimated to be 30-70% (5-7). Differences in phenotypic parameters that are used and age ranges of study participants may well contribute to the variation in the reported heritability. As summarized by Lewis et al. (8), genome-wide association studies (GWAS) of hearing status in adults and genetic analyses of families with dominantly inherited post-lingual onset HL indicate that both common variants and rare variants contribute to adult-onset HL with a small and large effect size, respectively. Such variants may or may not affect genes that are already known to function in the auditory pathway.

Previously, we identified a 12.4-Mb locus for adult-onset HL on chromosome 6 (p24.1-22.3): DFNA21 (9, 10). However, the underlying pathogenic variant in the studied family (W97-056) remained elusive. Here, we present the identification of an in-frame deletion (c.1696_1707del; NM_014722.3) in *RIPOR2* to underlie autosomal dominant nonsyndromic HL (adNSHL) in this family and in 11 additional (large) families of Dutch origin. The allele frequency (AF) of this variant suggests that it potentially explains adult-onset HL in thousands of individuals in the Netherlands and Northwest Europe.

## RESULTS

### Exome sequencing revealed an in-frame deletion in *RIPOR2*

To identify the genetic defect underlying the HL in family W97-056 (Figure 1), exome sequencing was performed in three affected family members (III:22, IV:20 and IV:25) which revealed one shared variants (AF ≤0.5%) that met the pathogenicity criteria and an in-frame deletion (Supplemental Table 1). A *SPATS1* variant (c.419G>A; p.(Gly140Glu); NM_145026.3), did not completely segregate with HL within the family as seven out of 23 affected subjects did not harbor the variant (Supplemental Figure 1). Also, *SPATS1* expression was not detected in the mammalian cochlea (11, 12) and SPATS1 function has only been related to spermatogenesis (13). Therefore, this variant was deemed non-causative. The in-frame deletion was present in exon 14 of *RIPOR2* (c.1696_1707del; p.(Gln566_Lys569del); NM_014722.3; Chr6:g.24,843,303_24,843,314del; rs760676508). It affects a highly conserved protein region of RIPOR2 (Rho family-interacting cell polarization regulator 2) which is present in all RIPOR2 isoforms (Supplemental Figure 2). *RIPOR2* has previously been associated with recessively inherited early-onset hearing loss and is positioned 0.9 Mb centromeric of the *DFNA21* locus (10, 14). No copy number variants (CNVs) were detected that were shared by all three subjects.

**Figure 1.**
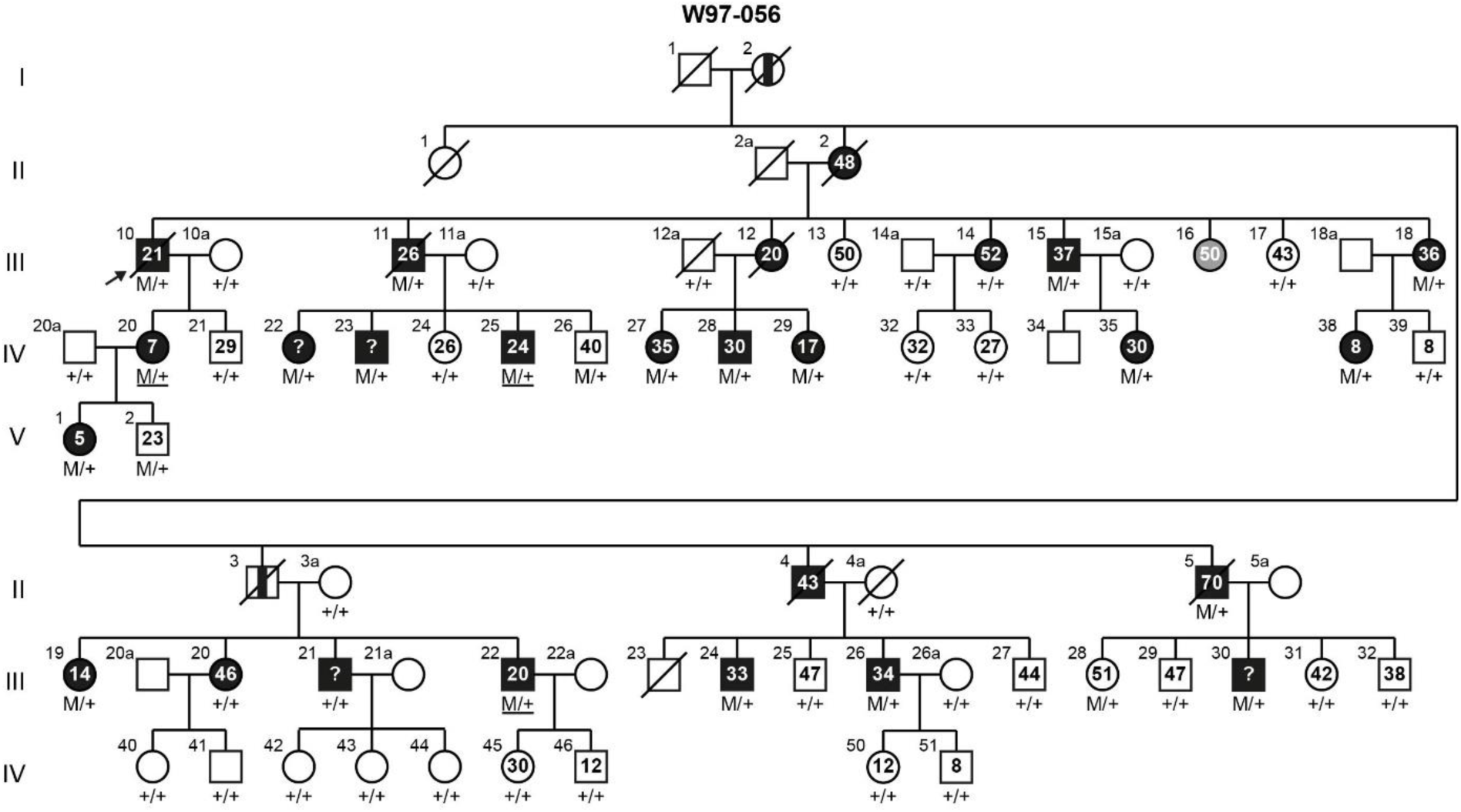
Pedigree of family W97-056 and segregation of *RIPOR2* variant c.1696_1707del. For affected and unaffected family members, the age of onset of hearing loss or the age at the most recent audiometric evaluation are indicated in the pedigree symbols, respectively. Subjects who did not report an age of onset are indicated with a question mark. The index case is marked by an arrow. Exome sequncing was performed in subjects with an underlined genotype. Subjects determined to be affected by heteroanamnesis are indicated with a vertical black bar. The subject marked in grey is diagnosed with intellectual disability and excluded from further participation in this study. Subject identifiers correspond to those in de Brouwer et al., 2005. M, c.1696_1707del; +, wildtype.

Segregation analysis identified the *RIPOR2* variant in 20 of 23 affected subjects of family W97-056 (Figure 1). The variant was not found in subjects III:14, III:20, and III:21; a recombination event in subject III:14 previously delimited the centromeric border of the *DFNA21* locus (10). The *RIPOR2* c.1696_1707del variant was also found in three unaffected family members (V:2, age 23 years; IV:26, age 40 years and III:28, age 51 years). The strong association of the *RIPOR2* variant with HL in this family urged us to further address this and other variants in *RIPOR2* in families with (adult-onset) HL.

### The *RIPOR2* variant c.1696_1707del associates with adNSHL in eleven additional families

An exome sequencing dataset of 1,544 index cases with (presumed) hereditary HL was evaluated for rare *RIPOR2* variants. In these cases, (likely) pathogenic variants in known deafness genes were previously addressed in a clinical diagnostic setting. The c.1696_1707del variant was identified in 10 index cases, all diagnosed with adNSHL (Figure 2). Analysis of a dataset obtained through Molecular Inversion Probe (MIP) sequencing of 89 HL-associated genes in 64 index cases with (presumed) adNSHL revealed another subject (V:1, W08-1421; Figure 2) with this variant. No other rare *RIPOR2* variants (AF ≤0.5%) that met the variant filtering criteria were identified.

**Figure 2.**
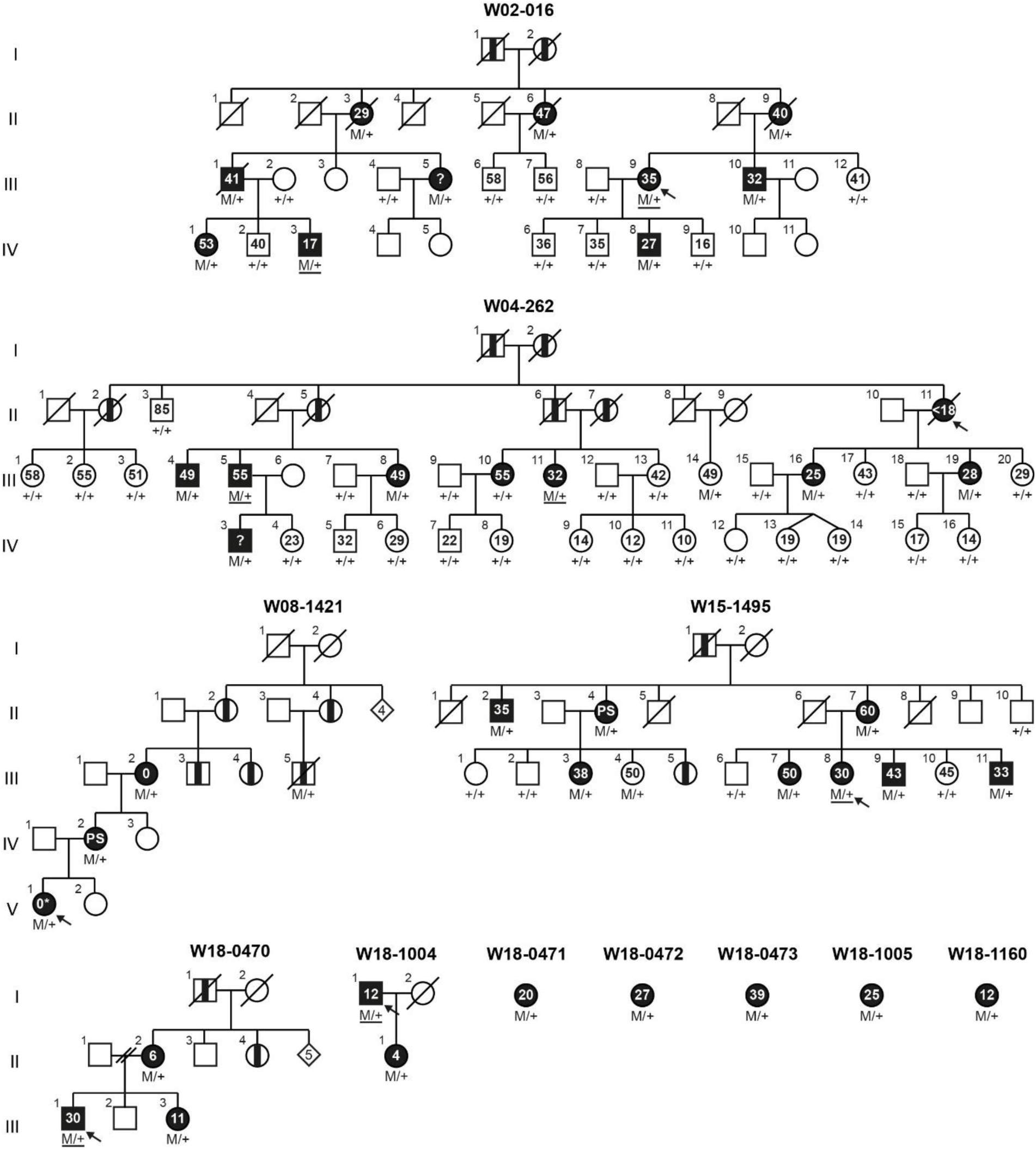
Family pedigrees and segregation of *RIPOR2* variant c.1696_1707del. For affected and unaffected family members, the age of onset of hearing loss or the age at the most recent audiometric evaluation are indicated in de pedigree symbols, respectively. Subjects who did not report an age of onset are indicated with a question mark. Index cases are marked by arrows. Exome sequencing was performed in subjects with an underlined genotype. Subjects determined to be affected by heteroanamnesis are indicated with a vertical black bar. Based on the information provided in the questionnaires, an autosomal dominant inheritance pattern of hearing loss is likely for each of the single cases. M, c.1696_1707del; +, wildtype; PS, primary school.

For six of the 11 index cases with the c.1696_1707del *RIPOR2* variant, family members were included in the study and segregation analysis was performed (Figure 2). The variant was detected in 39 of 40 affected subjects, but not in subject III:10 of family W04-262. As observed in family W97-056, the *RIPOR2* variant was also found in unaffected subjects namely III:14 of family W04-262 and III:4 of family W15-1495, aged 49 and 50 years respectively.

For all 11 index cases, targeted reanalysis of sequencing data for known adNSHL-associated genes (4) was performed to reveal other (likely) pathogenic variants. No rare variants were identified that both segregated with HL in the family and were classified as (likely) pathogenic in ClinVar (Supplemental Table 2).

### The *RIPOR2* c.1696_1707del variant does not affect transcript splicing

Splicing prediction algorithms (15) predicted that the deletion in exon 14 of *RIPOR2* may lead to the disruption of several exonic splicing enhancers. The loss of these could potentially induce alternative splicing events leading to premature truncation of protein synthesis. Therefore, RT-PCR was performed on RNA isolated from lymphoblastoid cells derived from patients and controls. No indications were obtained for an effect of the deletion on *RIPOR2* transcript splicing (Supplemental Figure 3).

### The *RIPOR2* c.1696_1707del variant is derived from a common ancestor

The presence of an identical *RIPOR2* variant in twelve families of Dutch origin is suggestive for a common ancestor. Variable Number of Tandem Repeats (VNTR) marker analysis was performed to determine whether a haplotype of the chromosomal region flanking the variant was shared by the different families. Indeed, a shared haplotype of ∼1.0 Mb, delimited by markers D6S2439 and D6S1281, was found in the seven families for which segregation analysis of the marker alleles could be performed (Supplemental Figure 4). This haplotype was also potentially shared by the five single cases. For marker D6S1545, a different CA-repeat length was determined on the variant-carrying allele of family W18-0470 whereas the alleles of two more centromeric markers where still shared. Since a rare event that caused a repeat length change of the D6S1545 allele may have occurred, this marker locus was still considered to be part of the shared haplotype.

To further refine the shared haplotype, we extracted homozygous SNPs present in the region between D6S2439 and D6S1281 from the exome sequencing datasets. Subsequently, homozygous SNP genotypes were compared between all index cases and discordant alleles were seen for SNP rs6901322 (Chr6: 24,583,804) that is located between D6S2439 and D6S1554. This SNP was found in homozygous state in subject IV:20 (W97-056), but was absent in the index cases of families W02-016 (III:9) and W18-0473. Based on these results, the shared haplotype is delimited by SNP rs6901322 at the telomeric side and comprises a region of 0.713 Mb. Genome sequencing in two members of family W97-056 (IV:25 and III:22) excluded potentially causative CNVs or other structural variants that are present within the shared chromosomal region.

### Clinical evaluation of individuals with the c.1696_1707del variant and phenocopies

To characterize the HL associated with the c.1696_1707del *RIPOR2* variant, 200 affected and unaffected subjects from seven families and five single index cases were evaluated between 1997 and 2018. The *RIPOR2* variant was found to be present in 64 of the 200 subjects. Detailed clinical data per individual are provided in Supplemental Table 3.

The mean reported age of onset is 30.6 years (standard deviation 14.9 years) with a wide range from congenital to 70 years (Supplemental Figure 5). Evaluation of audiometric data showed that subjects with the *RIPOR2* variant have progressive sensorineural HL, ranging from mild to profound, with variable audiometric configurations (Figure 3, Supplemental Figure 6). In order to distinguish audiometric patterns, a k-means clustering algorithm, independent of subject age, was applied on the latest audiogram of each subject. This unbiased approach yielded four audiometric patterns, each with a distinct audiometric configuration (Figure 4). Asymmetry of HL was seen in 16 cases. Inter-aural differences in progression of HL were also seen (Figure 3C). For three subjects (III:30 of family W97-056, II:11 of family W04-262 and V:1 of W08-1421), an explanation for asymmetry was noted (Supplemental Table 3).

**Figure 3.**
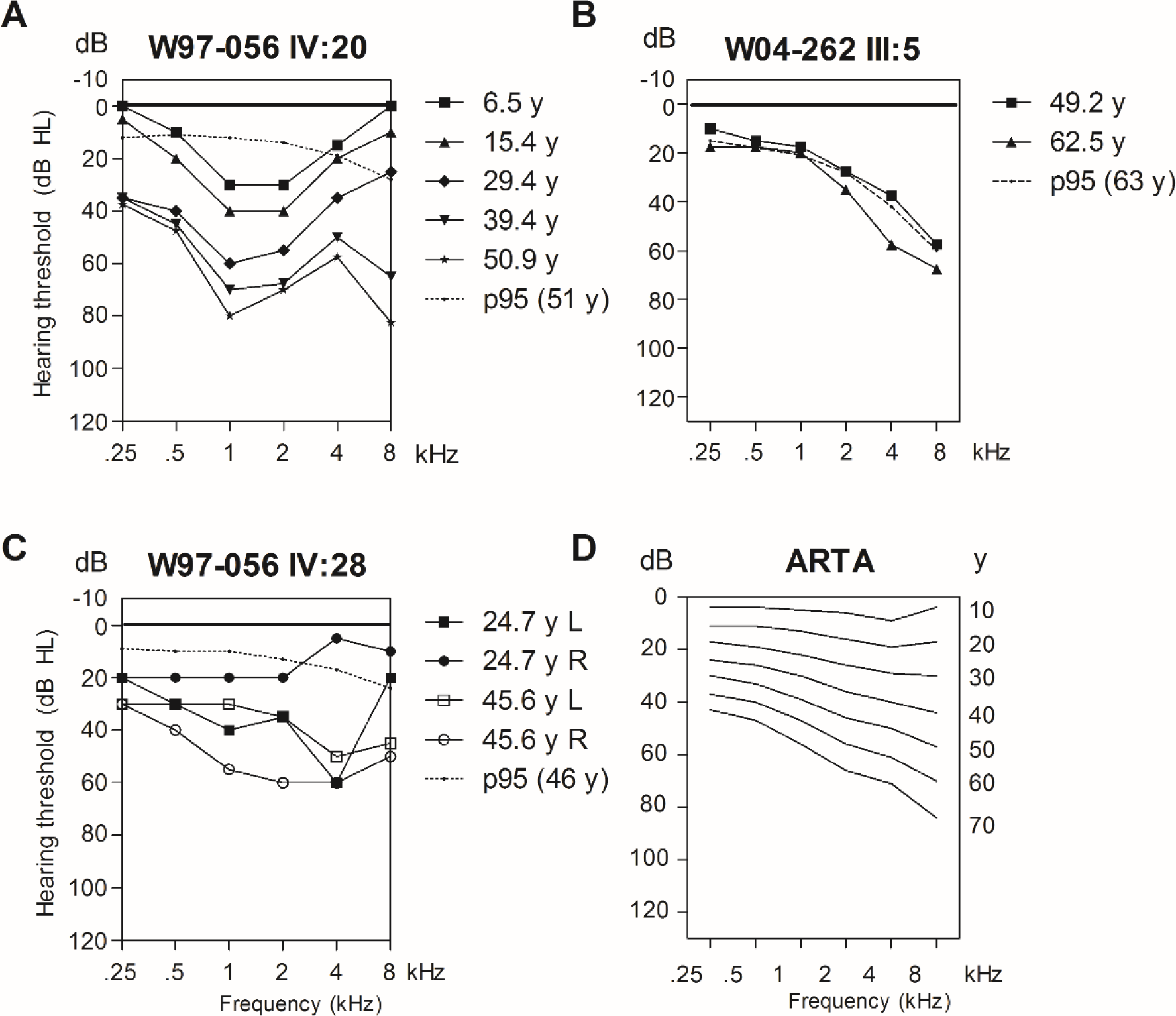
Selection of audiograms and the ARTA. **A-C.** Air conduction thresholds of three selected individuals with the c.1696_1707del *RIPOR2* variant are depicted. For the individuals in panels A and B, hearing loss was symmetric and the average of left and right ear thresholds are depicted. For the individual in panel C, hearing loss was asymmetric and the thresholds for both right and left ears are depicted.. The p95 values are matched to the individuals’ sex and age at most recent audiometry, according to the ISO 7029:2017 standard. **D.** Age Related Typical Audiogram (ARTA), cross-sectional linear regression analysis of last visit audiograms of affected subjects with the c.1696_1707del *RIPOR2* variant. y, age in years; R, right; L, left; dB HL, decibel hearing level; kHz; kiloHertz

**Figure 4.**
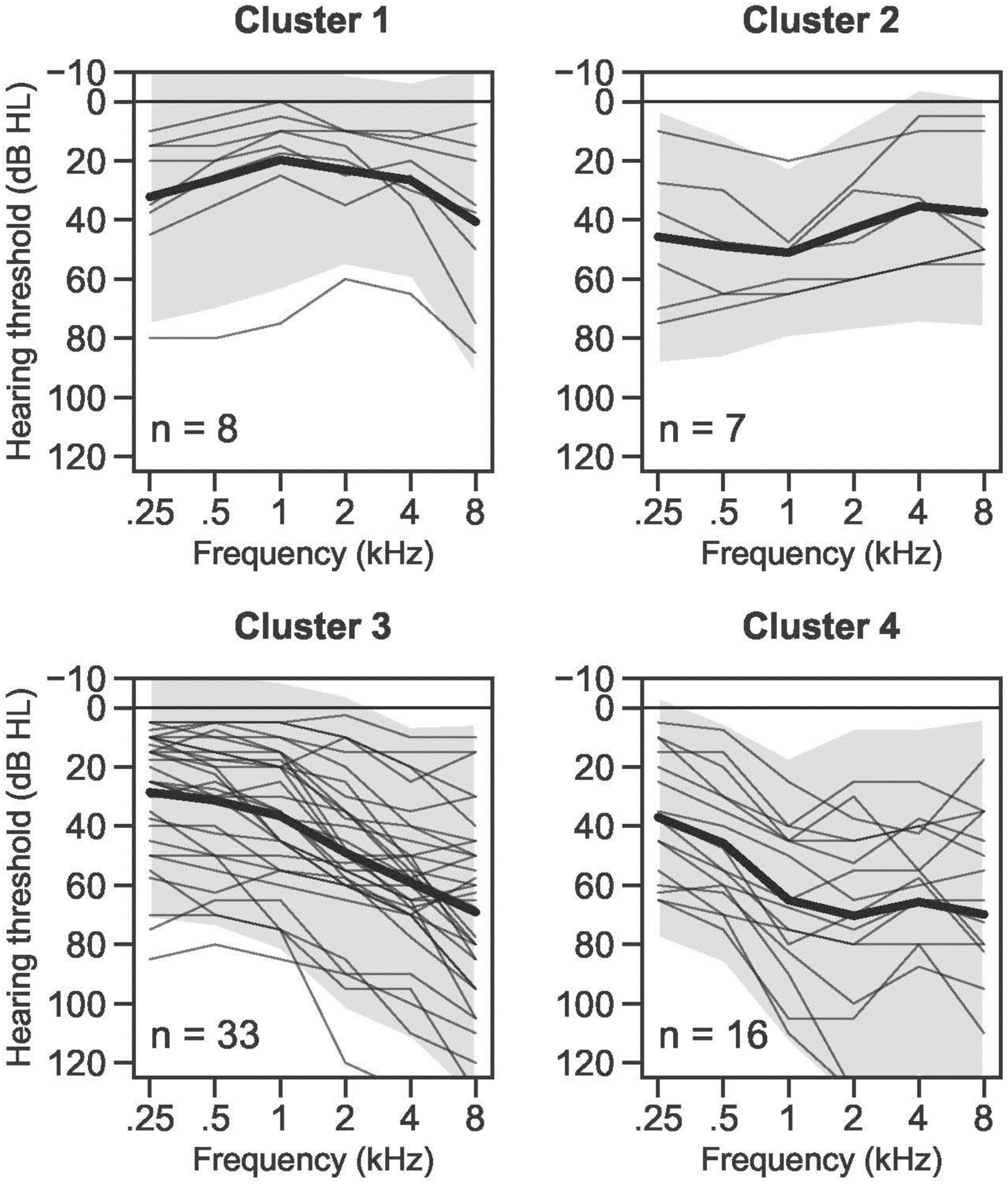
Four audiometric patterns of *RIPOR2*-associated hearing loss. Air conduction thresholds of all subjects were analysed with a k-means clustering protocol. The thick black lines depict the average of each cluster, the transparent grey areas represent the ±2 standard deviations. Cluster 1: mild hearing loss (average (PTA_0.5-4 kHz_)) 23 dB hearing level (HL) with an inverse U-shape audiogram. Cluster 2: moderate hearing loss (average 48 dB HL), with relatively worse hearing in the lower frequencies. Cluster 3: moderate (average 39 dB HL) high-frequency hearing loss with a gently down sloping audiogram configuration (average of 28 dB HL difference between the mean of 0.5-1 and 4-8 kHz). Cluster 4: moderate (average 60 dB HL), mid-frequency hearing loss with a U-shape audiogram, individual audiometry (Supplemental Figure 6) shows relatively faster deterioration of higher frequencies later in life, for example W97-056 IV:20. dB HL, decibel hearing level; kHz, kiloHertz.

Longitudinal analysis of HL in individual subjects revealed a large variation in progression of HL between subjects (Supplemental Table 3). We could not identify a specific pattern, such as a certain progression (in dB/y) in certain decades. There was a median progression of 1.2 dB/y (range 0.5-2.7 dB/y), for the frequencies 0.5-4 kHz. Cross-sectional linear regression was applied to calculate an Age Related Typical Audiogram (ARTA) (Figure 3D). Progression ranged from 0.7 dB/y (0.25 kHz) to 1.3 dB/y (8 kHz). Progression of HL was significant for all frequencies (*F*-test, p-value: <0.0001).

Speech reception thresholds were generally lower than, or comparable to, pure tone average at 0.5-2 kHz (PTA_0.5-2kHz_) (Supplemental Table 3). This indicates absence of retrocochlear pathology and is in line with normal results of click-evoked auditory brainstem response (ABR) in four subjects (Supplemental Table 3). CT and/or MRI of the bilateral temporal bones and cerebellopontine angle in six subjects revealed normal inner and middle ear anatomy (Supplemental Table 3).

Four of 64 subjects with the *RIPOR2* variant had vestibular complaints. Subjects III:1, III:9, IV:1 (W02-016) and the index cases of family W18-0472 reported infrequent vertigo attacks, complaints after cochlear implant surgery, a diagnosis of benign paroxysmal positional vertigo and migrainous vertigo, respectively (Supplemental Table 3). Vestibular testing was randomly performed in 9 subjects with the *RIPOR2* variant, aged 29 to 71 years, and included the abovementioned subjects III:1 and III:9 of family W02-016. No abnormalities were found, except for a mild hyporeflexia in subject III:1 of family W02-016 (Supplemental Table 4), which is appropriate for the subject’s age of 71 years. Based on these results, we conclude that c.1696_1707del *RIPOR2* is not associated with vestibular dysfunction. This is in line with the lack of vestibular dysfunction in *Ripor2*^-/-^ mice despite expression of the gene in the vestibular organ of wildtype mice (16). Also, humans with recessively inherited HL caused by a homozygous loss-of-function defect in *RIPOR2*, did not report balance problems, vertigo or dizziness but absence of a vestibular phenotype was not confirmed by objective vestibular testing (14).

Four subjects with HL who did not have the *RIPOR2* variant, are considered to be phenocopies (Supplemental Figure 7, Supplemental Table 5). For individuals III:14 and III:20 (W97-056) a possible explanation for their HL is a Ménière-like disease and heavy smoking (COPD Gold III), respectively (17, 18). Subject III:10 (W04-262) might have inherited a cause of HL associated with vestibular problems from her mother, who married into the family.

### Transcript levels of *RIPOR2* do not correlate with age of onset in affected subjects

We hypothesized that the variability in age of onset of the HL associated with the c.1696_1707del *RIPOR2* variant might be explained by differences in expression levels of the wildtype allele. Alternatively, variants in *cis-*regulatory elements of the affected allele more distantly located from *RIPOR2*, could influence expression levels of the mutant allele and might thereby modulate the age of onset. To test these hypotheses, allele-specific transcript levels of *RIPOR2* were determined in peripheral blood cells of 33 subjects using quantitative RT-PCR. Subjects were divided in three groups based on self-reported age of onset: <20 years (n=7), 20-39 years (n=15) and ≥40 years (n=6). No significant differences were observed between the different subject groups, neither for the wildtype or c.1696_1707del variant *RIPOR2* alleles nor for total *RIPOR2* transcript levels (Supplemental Figure 8). Also, no difference was observed between the ratios of *RIPOR2* mutant to wildtype relative transcript levels. A small difference was observed in total *RIPOR2* transcript levels between subjects with an early onset of HL and controls (p=0.0241). This could suggest a trend between low expression levels and an early onset of HL, however, considering the overall variability in transcript levels it is more likely that other factors play a role. A larger sample size would be required to confirm or negate the observed trend.

### The in-frame deletion in *Ripor2* prevents correct localization of the protein in mouse cochlear hair cells

Previous studies have shown that RIPOR2 is specifically localized to the base of the stereocilia in mouse cochlear hair cells (19). RIPOR2 is highly conserved between mouse and human (87% amino acid identity). To study whether the localization of mouse RIPOR2 with a deletion of the orthologous four amino acid residues (p.584_587del) is altered, plasmids encoding wildtype- or mutant-RIPOR2 were injectoporated into cochlear outer hair cells of wildtype mice (P2). Interestingly, mutant-RIPOR2 was detected in the stereocilia but in none of the 12 evaluated cells it was retained at the stereocilia base where the wildtype protein was found to be localized in all 11 evaluated cells (Figure 5A). Morphology of the stereocilia was not significantly affected two days after injectoporation of the mutant *Ripor2* construct, suggesting the mutant protein did not visibly affect the stereocilia structure in the short term.

**Figure 5.**
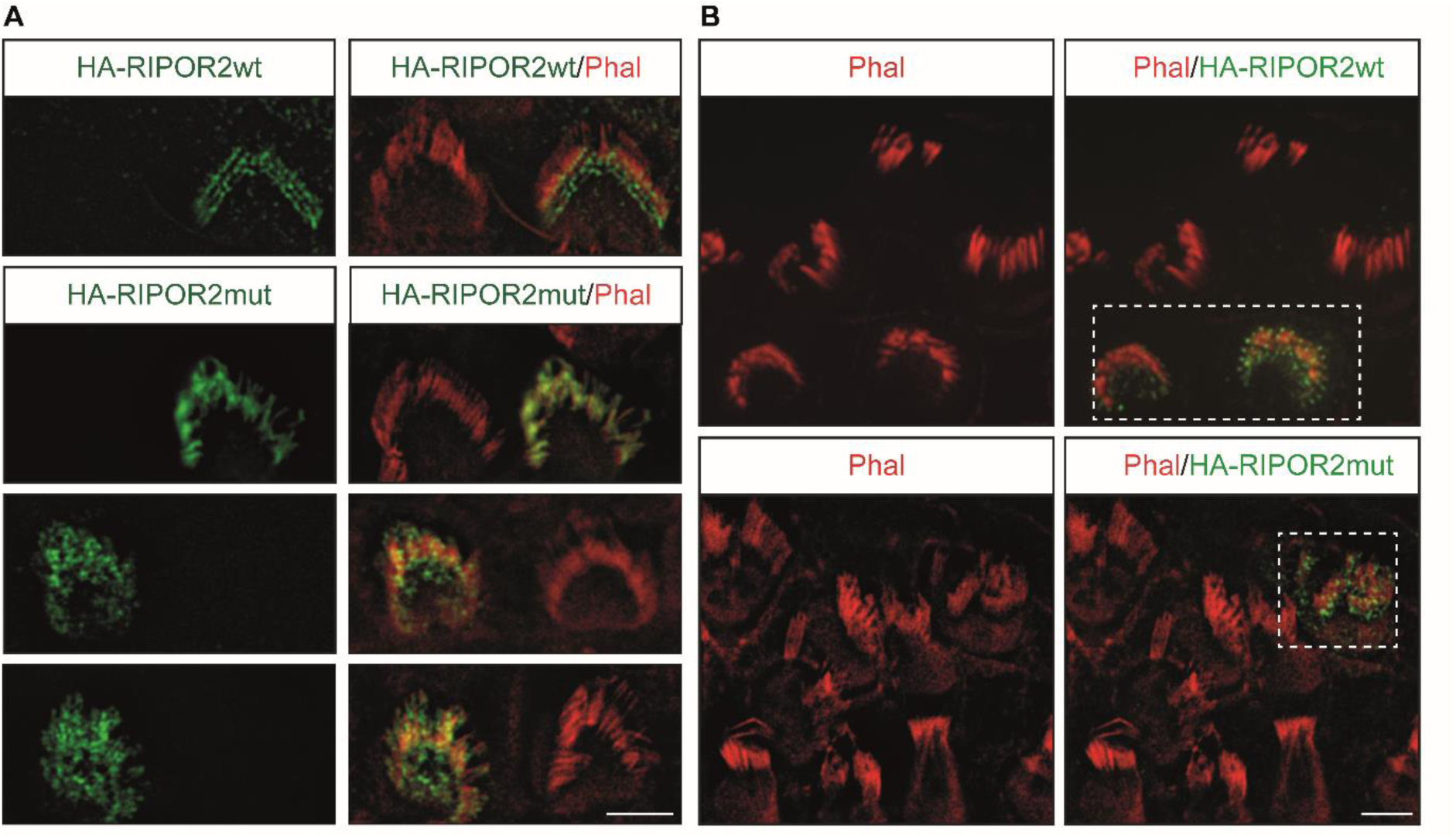
Functionality of mutant RIPOR2 is altered in mouse cochlear outer hair cells. **A.** Mutant RIPOR2 differed in localization from wildtype RIPOR2 in mouse cochlear outer hair cells. Outer hair cells of wildtype mice were injectoporated at P2 to express murine N-terminally HA-tagged wildtype RIPOR2 (RIPOR2wt) or mutant RIPOR2 (RIPOR2mut). Expression was evaluated after two days by immunohistochemistry and three representative images of cells expressing the mutant RIPOR2 are provided. Eleven cells expressing the wildtype construct and 12 cells expressing the mutant construct were evaluated. **B.** Mutant RIPOR2 did not rescue stereocilia defects in Ripor2-deficient hair cells. Cochlear explants of RIPOR2-deficient mice were prepared at P2 and injectoporated with constructs RIPOR2wt or RIPOR2mut. After culturing for two days, five out of six cells expressing the wildtype RIPOR2 construct demonstrated rescued hair bundle morphology but none of the 13 cells expressing the mutant RIPOR2 construct. Cells expressing the constructs are boxed. HA-tagged protein was stained in green, stereocilia were stained using phalloidin (phal) conjugated with Alex Fluor 568 (red). Scale bar represents 5 μm.

Potentially, the variant affects interactions of RIPOR2 that are essential for its localization. The four deleted amino acids are predicted to be part of a disorganized coiled coil structure (predicted using KMAD (20)). Coiled coil regions are indicated to mediate protein-protein interactions, which supports the hypothesis that the variant affects RIPOR2 protein interactions (21). Co-immunoprecipitation (Co-IP) assays demonstrated that both the dimerization ability (Supplemental Figure 9A) and the interaction with RHOC (Supplemental Figure 9B) of mutant RIPOR2 are intact.

### Mutant RIPOR2 cannot rescue morphological defects in outer hair cells from *Ripor2* knockout mice

In *Ripor2* knockout mice, morphological defects were previously observed in hair cells, which included bundle polarity, cohesion and length of stereocilia (19). After injectoporation of the *Ripor2* mutant construct into outer hair cells of these mice, these defects could not be rescued in any of the 13 cells expressing mutant-RIPOR2. The typical V-shaped hair bundle was not formed, in contrast to the rescue effect observed in five out of six cells expressing wildtype-RIPOR2 (Figure 5B). This, together with the aberrant localization of mutant-RIPOR2, confirms an effect of the 4-amino acid deletion on RIPOR2 function in outer hair cells.

## DISCUSSION

This study identified an in-frame 12 nucleotide deletion in *RIPOR2* as a prevalent and highly penetrant genetic factor for adult-onset HL in the Netherlands and beyond. HL associated with the deletion is highly variable in age of onset and audiometric characteristics. Our study exemplifies that an increasing contribution of environmental factors and of low-penetrance genetic factors to the hearing ability during life, complicates the identification of highly penetrant genetic factors in adult-onset HL. This is best illustrated by family W97-056 in which the linkage interval was falsely delimited by a phenocopy.

The *RIPOR2* variant was significantly enriched in an in-house dataset, previously coined “SE-NL” (Southeast Netherlands) with exomes of 22,952 unrelated individuals with unknown hearing abilities (22). Eighteen individuals were heterozygotes for the variant (AF 0.0392%), as compared to 8 of 56,352 individuals (AF 0.0071%) and 5 of 32,287 individuals (AF 0.0077%) of non-Finnish European (nFE) descent in the gnomAD exome database v2.1.1 and gnomAD genome database v3, respectively. As the variant was indicated to be inherited from a common ancestor, this individual might well by of Dutch origin or of neighboring regions.

Several lines of evidence indicate the association of the c.1696_1707del *RIPOR2* variant with HL. Firstly, the deletion affects four highly conserved amino acids of RIPOR2, which is known to have a crucial role in murine and zebrafish hair cell development, function, and maintenance (14, 16, 19). *Ripor2* knockout mice are already found to be deaf at four weeks of age due to impaired mechanotransduction (19). Also, knockdown of *ripor2* in zebrafish induced loss of hair cells, and consequently profound hearing loss (14). Secondly, aberrant localization of the mutant RIPOR2 in early postnatal mouse hair cells, *ex vivo*, and failure to rescue the stereocilia defects of *Ripor2* knockout mice indicate a functional effect of the variant. Thirdly, neither other rare potentially causative variants in protein coding regions and splice sites of the shared haplotype region, nor structural variants affecting this region were revealed in exome or genome sequencing.

RIPOR2 is localized at the taper region of the mechanically sensitive stereocilia of murine hair cells (14, 16, 19) where it is organized in a ring-like fashion (19). The latter is thought to be achieved by homo-oligomerization in a head-to-head and tail-to-tail manner, regulated by RHOC (19). The oligomerization is essential for the structure of the taper region and for the morphology of the hair bundle as a whole, but the precise molecular mechanism is still elusive. The taper region is the specialized basal part of stereocilia that allows their deflection upon mechanical stimulation (23). CLIC5, PTPRQ, MYO6, TPRN, RDX, GRXCR2, and RIPOR2 are described to concentrate and co-function in the taper region and to be crucial for its structure and/or for hair bundle development and maintenance in mice (19, 24-29). Direct interactions of these proteins are indicated, e.g., of CLIC5, RDX and TPRN, but not RIPOR2 (19, 26). Also, interdependence for their concentration in the taper region was observed (19, 25, 26). In RIPOR2-deficient hair cells, for example, TPRN is no longer concentrated at the stereociliary base (19, 24). Depletion of TPRN in *Tprn* knock-out mice leads to functional as well as (slowly) progressive morphological abnormalities of the stereocilia bundle (24).

Based on the above described molecular structure of the stereociliary taper, we hypothesize that p.(Gln566_Lys569)del RIPOR2 affects this taper region and thereby the durability of the hair bundle, potentially via an effect on TPRN. Additionally or alternatively, the *RIPOR2* variant might affect the amount of the RIPOR2-interaction partner MYH9 in stereocilia, as well as the abundance of phosphorylated MYH9 and acetylated α-tubulin in the kinocilia, as these proteins are reduced in RIPOR2-deficient mice (16). Interestingly, *MYH9* defects in humans are also associated with progressive HL (30).

In light of developing therapeutic strategies, it is essential to determine whether the *RIPOR2* variant has a loss-of-function, a dominant negative or toxic gain-of-function effect. A haploinsufficiency effect of the variant seems to be the least plausible, as a loss-of-function *RIPOR2* variant in the heterozygous state was not indicated to be associated with HL (14). Also, heterozygous *Ripor2* knockout mice displayed no significant hearing loss at four weeks (19) and two months of age (Zhao, unpublished data). A dominant-negative effect of the p.(Gln566_Lys569del) variant cannot be excluded as an interaction between the mutant- and wildtype-RIPOR2 was detected in Co-IP assays. However, a strong dominant negative effect would be expected to result in early-onset HL, comparable to that associated with the homozygous loss-of-function variant (14). Therefore, we hypothesize that the variant has a toxic gain-of-function effect.

RIPOR2 is expressed in a wide-range of tissues and cell types (19). It is a known inhibitor of the small G-protein RHOA in neutrophils and T lymphocytes, where it regulates migration of these cells (31). Additionally, *RIPOR2* is upregulated during muscle cell differentiation and induces the formation of filopodia (32). We did not observe an effect of the four amino acid-deletion on filopodia formation (de Bruijn, unpublished data) which is in line with the fact that the deleted residues are not part of the RHOA-interaction domain (32). This might, at least in part, explain that the *RIPOR2* variant leads to HL only. The variant could affect a cochlear-specific protein interaction that determines RIPOR2 localization in the hair bundle. Furthermore, in tissues other than the inner ear loss of RIPOR2 function might be compensated by RIPOR1 and RIPOR3 which are described to have redundant functions (33, 34). Indeed, RNA levels of both *RIPOR1* and *RIPOR3* are low in hair cells (https://umgear.org/).

The audiometric phenotype and age of onset of HL associated with c.1696_1707del *RIPOR2* displayed variation. Such intrafamilial phenotypic variation has also been reported for defects in several of the genes that can be associated with adult-onset adNSHL, e.g. *EYA4, MYO6* and *POU4F3*, and remains unexplained (35-37). With a k-means cluster analysis, four distinct audiometric clusters could be distinguished. It is possible that subjects, due to increasing age, may go from one cluster to another cluster, which is not captured by the k-means clustering algorithm, since no longitudinal data are used. As no clear patterns of age of onset or audiometric configurations were observed within families or family branches with the *RIPOR2* variant, the phenotypic variability might well result from an interplay between environmental and genetic modifying factors. We have addressed differences in transcript levels of both wildtype and mutant *RIPOR2* alleles as potential modifiers of age of onset but no clear correlations were observed. As the analysis was performed on RNA extracted from peripheral blood, we cannot exclude that *RIPOR2* mRNA levels determined by cochlear-specific *cis* or *trans* regulatory elements modify the onset of HL. Other candidate genetic modifiers are variants in the genes that encode proteins of the indicated complex of the stereocilia taper. As the taper region is thought to be essential for anchoring the mechanosensory stereocilia, noise exposure is an obvious candidate environmental modifying factor. Fourteen subjects with the *RIPOR2* variant reported noise exposure. However, we could not correlate onset or strong progression of HL with a preceding significant noise exposure.

The c.1696_1707del *RIPOR2* variant was only reported in non-Finnish Europeans, with the exception of a single individual of African origin (gnomAD v3 genomes). Assuming that the AF of 0.0392% determined in the SE-NL cohort is comparable throughout the Netherlands, the c.1696_1707del *RIPOR2* variant is estimated to be present in more than 13,000 individuals who are therefore at risk to develop HL or have developed HL already due to this variant. About 30,000 additional individuals can be calculated to be at risk, based on the AF of 0.0096% of the variant in Northwest Europe (gnomAD v2.1.1) with ∼156 million inhabitants (United Nations Population Division estimates, 2019). This large number of individuals at risk to develop HL due to the c.1696_1707del *RIPOR2* variant illustrates the need to gain broader estimates of the penetrance of the variant which was ∼90% at the age of 50 years in the studied families. However, this calculated penetrance cannot be excluded to be biased because these families were included based on index cases with HL. Further insight in the age-related penetrance of c.1696_1707del *RIPOR2* will pave the way for the identification of modifying factors which may convey handles for prevention.

In conclusion, we demonstrate that an adult-onset type of HL (DFNA21) is relatively common and associated with a ‘mild’ variant in *RIPOR2*. Potentially, thousands of individuals in the Netherlands and beyond are at risk to develop HL. More such variants might well wait to be ‘unmasked’ as (population-specific) frequent and highly penetrant causes of adult-onset HL. Because of the large number of subjects estimated to be at risk for HL due to the c.1696_1707del *RIPOR2* variant, it is an attractive target for the development of a genetic therapy. The great progress that is being made for this in hearing disorders is promising (38).

## METHODS

### DNA sequencing

Genomic DNA was isolated from peripheral blood lymphocytes following standard procedures. Subsequently, exome enrichment was performed using the Agilent SureSelect Human All Exome V5 kit according to the manufacturer’s protocols. Exome sequencing was performed on an Illumina HiSeq system by BGI Europe (Copenhagen, Denmark). Read mapping along the hg19 reference genome (GRCh37/hg19) and variant calling were performed using BWA V.0.78 (39) and GATK HaplotypeCaller V.3.3 (40). A coverage of >20 reads was reached for 85.1% to 97.8% of the enriched regions. For variant annotation an in-house developed annotation and variant evaluation pipeline was used. For sequencing data of family W97-056, CNV detection was performed using CoNIFER V.0.2.2 (41). Genome sequencing was performed by BGI (Hong Kong, China) on a BGISeq500 using a 2x 100 bp paired end module, with a minimal median coverage per genome of 30-fold. Structural variants were called using Manta V.1.1.0 (42) and CNVs using Control-FREEC (43). Variants were validated and visualized using the IGV Software (V.2.4) (44).

In the index case of family W08-1421, targeted DNA sequencing was performed using MIP sequencing (45). MIPs were designed covering exons and exon-intron boundaries of a panel of 89 HL genes (Supplemental Table 6). Sequencing and data analysis were performed as previously described (46). For each targeted region, an average coverage of 420 reads was obtained. A coverage of >20 reads was reached for 85.4% of the MIPs. Only those called variants were considered that had a quality-by-depth >200 and that were present in less than 10% of the samples that were analyzed in the same sequence run (n=150).

### Variant interpretation

For exome sequencing and MIP datasets, annotated variants were filtered based on a population AF of ≤0.5% in the gnomAD database V.2.1 (http://gnomad.broadinstitute.org), and our in-house exome database (∼15,000 alleles). Variants in coding and splice site regions (−14/+14 nucleotides) were analyzed. Interpretation of missense variants was performed using the in silico pathogenicity prediction tools CADD-PHRED (≥15) (47), SIFT (≤0.05) (48), PolyPhen-2 (≥0.450) (49) and MutationTaster (deleterious) (50). Variants were considered if a pathogenic effect was predicted by at least two different tools. Potential effects on splicing of missense and synonymous variants were evaluated using the algorithms embedded in the AlamutVisual software (V.2.10, Interactive Biosoftware). A change of ≥5% in splice site scores predicted by at least two algorithms was considered significant. For candidate variants, segregation analysis was performed by Sanger sequencing. PCR conditions are available upon request.

### Splicing assay

Potential effects on splicing of the *RIPOR2* variant were assessed using the splicing prediction tools via the AlamutVisual software as described above and using Human Splicing Finder V.3.1 to predict effects on exonic splicing enhancers and silencers (15). Predicted alternative splicing events were experimentally addressed by RT-PCR. For this, total RNA was isolated from patients’ cultured lymphoblastoid cells. For cDNA synthesis with the iScript cDNA synthesis kit (Bio-Rad), 1 μg RNA was used as input. RT-PCR was performed with primer combinations for exons 11-16 and 12-15 of *RIPOR2* (Supplemental Table 7), and followed by visualization of amplified products using agarose gel electrophoresis.

### VNTR marker analysis

Genotyping of VNTR markers was performed by genomic DNA amplification using touchdown PCR and analysis on an ABI Prism 3730 Genetic Analyzer (Applied Biosystems). Genomic positions of markers were determined using the UCSC genome browser (human genome assembly GRCh37/hg19). Alleles were assigned with the GeneMarker software (V.2.6.7, SoftGenetics) according to the manufacturer’s protocol.

### Clinical evaluation

Medical history was taken from all participants with special attention paid to acquired and noise-induced HL. Both affected and unaffected participants underwent general Ear Nose and Throat examinations, or this medical information was taken from previous examinations. Age of onset of HL was reported by subjects themselves. Only reports of a specific age of onset were used in calculations. The audiometric data in this study are described according to GENDEAF guidelines (51). Pure tone- and speech-audiometry and click-evoked ABR was performed in a sound-attenuated booth, according to current standards (International Organisation for Standardization; ISO 8253-1:2010, ISO 389-1, ISO 389-5 and ISO 389-6) (52). Individuals were considered affected when pure tone thresholds for at least three individual frequencies were below the frequency-specific 95^th^ percentile of age- and sex-specific thresholds (ISO7029:2017) for the best hearing ear. HL was considered asymmetric if pure tone audiometry showed a difference of more than 10 dB between both ears at two individual frequencies (51). Longitudinal (individual) progression of HL was calculated if there was a follow-up duration of at least 10 years, after onset of HL. The progression rate is defined as the mean increase PTA_0.5-4kHz_ in dB/year between first and last audiometry. For symmetric HL, the average of both ears was used to calculate progression; for asymmetric HL, the best-hearing ear at first audiometry was used. In case of profound HL at 0.5-4kHz at the latest audiometry, the most recent audiometry at which all thresholds at 0.5-4kHz could be measured, was selected. Cross-sectional linear regression analysis was applied on pure tone thresholds to calculate an ARTA (53), using Prism 6.0 software (GraphPad).

Vestibular function was assessed by electronystagmography, caloric irrigation testing, rotary chair stimulation and video head impulse tests, as described previously (54). Cervical and ocular vestibular-evoked myogenic potentials (cVEMP/oVEMP) were measured to assess saccular and utricular function, respectively (55, 56). When responses were seen at or below 100 dB Hearing Level during (air conducted) cVEMP testing, saccular function was considered to be present, otherwise absent (55). For (bone conducted) oVEMP stimulation, this normal value is ≤140 dB Force Level (56).

### Audiometric cluster analysis

A k-means clustering algorithm was applied on the last audiogram of affected subjects with the *RIPOR2* variant (57). Each audiogram was first normalized by subtracting the average hearing threshold across all frequencies from the data, preventing the algorithm to select clusters based on average hearing threshold, while retaining the overall shape of the audiogram. Next, the optimal number of clusters in the data was obtained by using the Elbow method (58); for a number of k=1 to k=15 clusters. The distortion, i.e. the sum of squared distances from each point to its assigned center, was determined for each value of k. The smallest number of k clusters with the lowest distortion (i.e. the elbow) was then taken as the optimal number of clusters. Finally, the audiograms of all the patients within each cluster were visualized and an average cluster prototype was obtained by averaging all audiograms within a cluster.

### Allele-specific expression analysis

Peripheral blood (2.5 ml) was collected in PAXgene Blood RNA tubes (BD Biosciences). RNA was isolated using the PAXgene blood RNA kit (Qiagen) following the manufacturer’s protocol. Subsequently, cDNA was prepared using the iScript cDNA synthesis kit (Bio-Rad) with 500 ng RNA. Allele-specific primer sets were designed and validated; the design was based on a forward primer that specifically hybridizes to either the mutant or wildtype *RIPOR2* alleles. Additionally, primers were designed for exons 3-4 of *RIPOR2*, as well as for exons 2-3 of the reference gene *GUSB* (NM_000181). Primer sequences are provided in Supplemental Table 7. All qPCR reaction mixtures were prepared with the GoTaq qPCR Master Mix (Promega) according to the manufacturer’s protocol. Amplifications were performed with the Applied Biosystem Fast 7900 System (Applied Biosystems). For all RNA samples, cDNA was synthesized twice, and all qPCR reactions were performed in duplicate. Relative gene expression levels, as compared to the internal reference gene *GUSB*, were determined with the ΔCt method (59). Statistical analyses were performed using a one-way ANOVA followed by Tukey’s multiple comparison test to test for significance between the groups.

### Injectoporation of *Ripor2*-constructs and immunostaining

The generation of *Ripor2*^*LacZ/LacZ*^ mice has been described previously (19). For *Ripor2* DNA construct generation, *Ripor2* cDNA (NM_029679.2, without exon 13) was amplified from a mouse-cochlear cDNA library and cloned into a pEGFP-N3-derived vector in which the EGFP was deleted. For injectoporation, the organ of Corti was isolated and placed in DMEM/F12 medium with 1.5 μg/ml ampicillin. Glass electrodes (∼2 μm diameter) were used to deliver the plasmid (500 ng/μl in Hank’s Balanced Salt Solution (HBSS)) to the sensory epithelium. A series of 3 pulses were applied at 1 sec intervals with a magnitude of 60V and duration of 15 msec (ECM 830 square wave electroporator; BTX). Two days after injectoporation, samples were fixed in the fixative containing 4% paraformaldehyde in HBSS for 20 min. Tissues were then washed in HBSS and blocked for 20 min at room temperature in HBSS containing 5% BSA, 1% goat serum and 0.5% Triton X-100, and then incubated overnight at 4°C with primary antibodies in HBSS containing 1% BSA and 0.1% Triton X-100. Tissues were washed in HBSS and incubated 2 hours at room temperature with secondary antibodies. Tissues were mounted in ProLong^®^ Antifade Reagents (ThermoFisher). Stacked images were then captured by fluorescence deconvolution microscope (Leica). Antibodies used were: anti-HA (mouse; 1:500; cat.#2367S; Cell Signaling), Alex Fluor 568-phalloidin (1:500; cat.#A12380; ThermoFisher) and Alexa Fluor 488 goat anti-mouse (1:1000; cat.#A11017; ThermoFisher).

### Immunoprecipitations and western blots

HEK293T cells were obtained from ATCC and were maintained in DMEM medium (ThermoFisher) supplemented with 10% heat-inactivated fetal bovine serum and 1% penicillin/streptomycin. Cells were grown at 37°C in a 5% CO_2_ humidified atmosphere. Immunoprecipitations and western blots were performed as described (19). Experiments were carried out at least 3 times to verify the reproducibility of the data. The following antibodies were used for the experiments: anti-HA (mouse; 1:500; cat.#2367S; Cell Signaling), anti-Myc (rabbit; 1:500; cat.#2278S; Cell Signaling), anti-Myc (mouse; 1:500; cat.#9E10; Santa Cruz); anti-GFP (mouse; 1:1000; cat.#SC-9996; Santa Cruz).

### Study approval

The study of human subjects was approved by the medical ethics committee of the Radboudumc (registration number: NL33648.091.10) and performed in accordance with the principles of the World Medical Association Declaration of Helsinki. Written informed consent was obtained from all participants or their legal representatives. All animal experiments were approved by the Institutional Animal Care and Use Committee of Indiana University School of Medicine (registration number 19075).

## AUTHOR CONTRIBUTIONS

SEdB and JJS co-designed the study, conducted experiments, analyzed data, wrote the manuscript and JJS performed subject evaluation. CL performed the experiments in mouse cochlear explants- and Co-IP-experiments and revised the manuscript. CPL and AJB analyzed the audiovestibular data and revised the manuscript. EdV discussed experimental design and critically read the manuscript. The DOOFNL consortium is a Dutch nationwide collaboration on hereditary hearing loss. Specifically for this study; DOOFNL-members RJCA, SGK, LJCR, SMM and JvdK contributed subjects with the *RIPOR2* variant for this study. Other members of the consortium contributed subjects to the cohort of index cases. JB and JO conducted and analyzed genetic analyses. WK and HGY analyzed WES-data and revised the manuscript. FPMC and SR, discussed experimental design and critically read the manuscript. HPMK performed clinical evaluations for members of family W97-056. BZ co-designed and supervised the studies in mouse cochlea and the Co-IP-experiments, and revised the manuscript. RJEP clinically evaluated family members, RJEP and HK co-designed the study, supervised the project and revised the manuscript. All authors read and approved the final manuscript.

## ACKNOWLEDGMENTS

The authors thank José van den Broek, Herman Kok, Lisanne Schenk, Daniëlle Manders, Karin Krommenhoek and Jacquelien Jilissen for their contributions to clinical evaluations and Saskia van der Velde-Visser for her contributions in EBV transformations and cell culture. We would like to thank Sita Vermeulen, Galuh Astuti and Hanka Venselaar for their contribution to statistical and bioinformatic analyses. This study was financially supported by a DCMN Radboudumc grant, by a grant of the Heinsius-Houbolt foundation and an NIH/NIDCD (R01 DC017147) grant. See Supplemental Acknowledgments for consortium details.

