## Supplementary material for "A *RIPOR2* in-frame deletion is a frequent and highly penetrant cause of adult-onset hearing loss": suppl figures and tables

**SUPPLEMENTAL FIGURES**

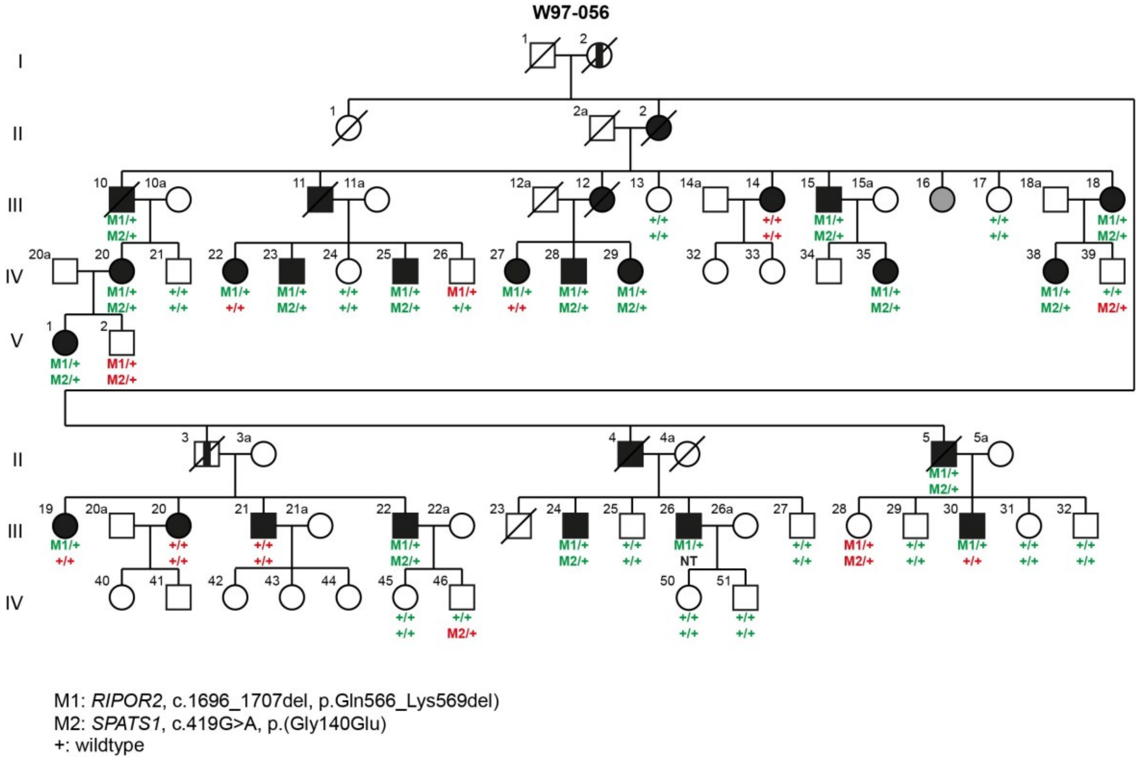


**Supplemental Figure 1. Segregation analysis of the *SPATS1* and *RIPOR2* variants in W97-056.** Subjects determined to be affected by heteroanamnesis are indicated with a vertical black line. The subject marked in grey is diagnosed with intellectual disability and excluded from further participation in this study. Subject identifiers correspond to those in de Brouwer et al., 2005 (10). Genotypes in green correspond to a co-occurrence of the variant and hearing impairment and those in red to lack of co-occurrence. NT, not tested.

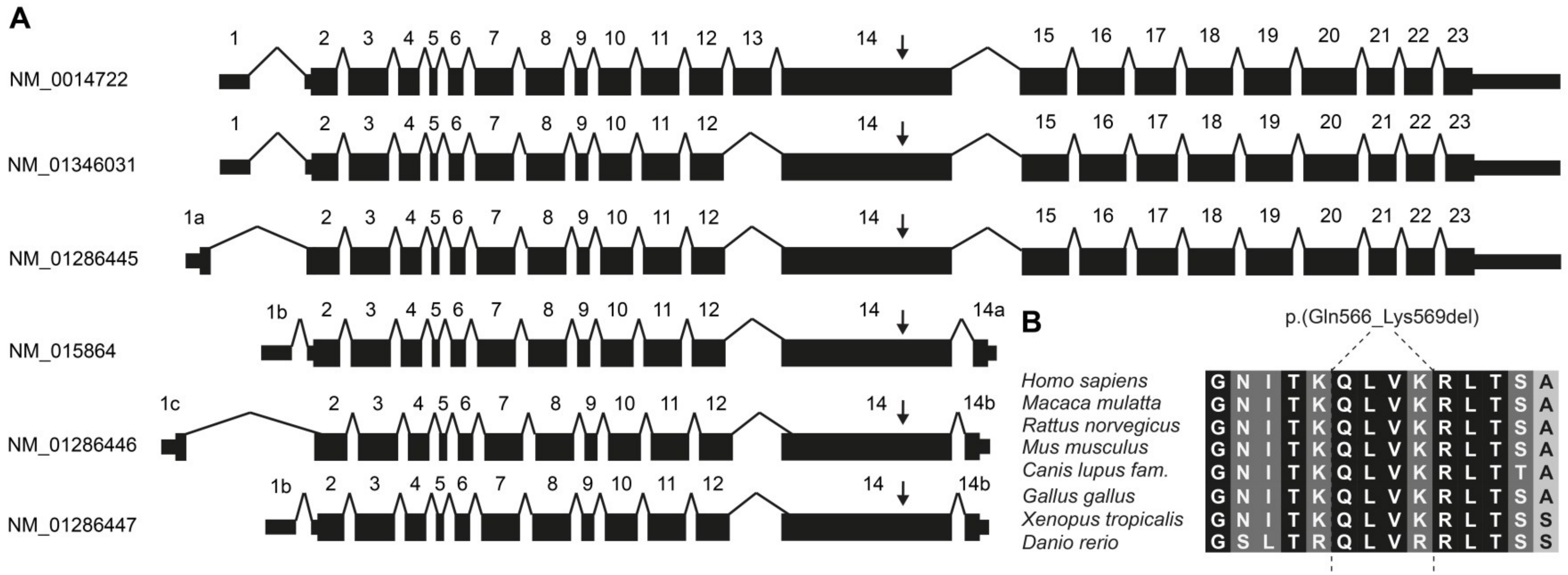


**Supplemental Figure 2. Schematic overview of major *RIPOR2* transcripts.** (A) Transcripts are extracted from the Ensembl genome browser. (B) Evolutionary conservation of the amino acids that are affected by the variant. Fully conserved amino acid residues are shown on a black background. A dark grey background marks chemical similarity of residues and a light grey background indicates chemical dissimilarity.

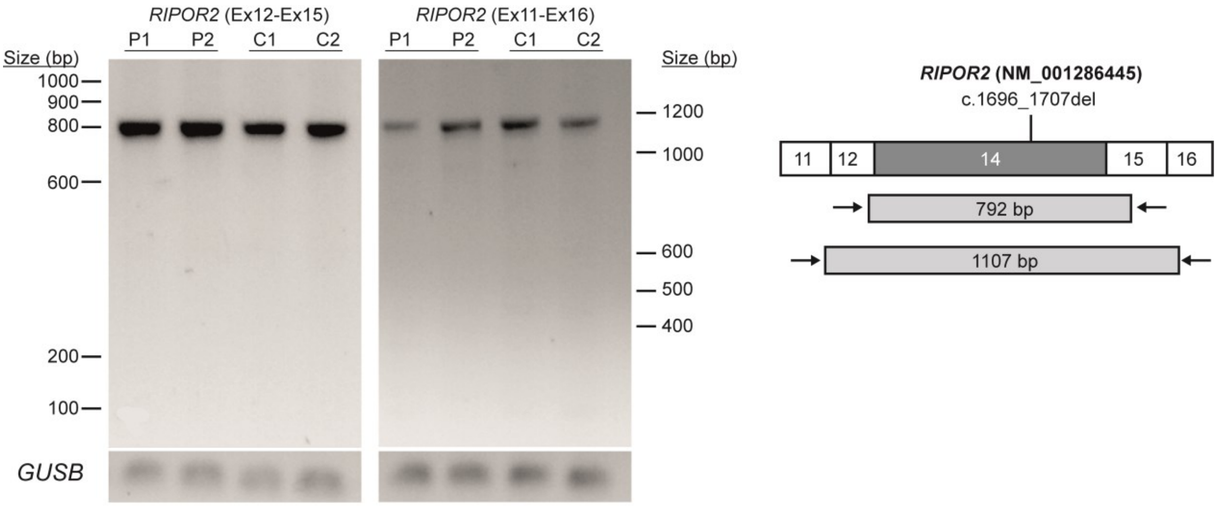


**Supplemental Figure 3. Analysis of *RIPOR2* transcript splicing**. RT-PCR using primer pairs for the exons flanking exon 14 in transcript NM_001286445.2 was performed to detect aberrant splicing potentially induced by the *RIPOR2* c.1696_1707del variant. The amplicon sizes for the expected normal splicing events are indicated in the right panel of the figure. RNA was isolated from lymphoblastoid cells from patients (P1 and P2; W97-056: IV:20 and III:26) or controls (C1 and C2) who did not have the *RIPOR2* variant. *GUSB* was used as cDNA input control.

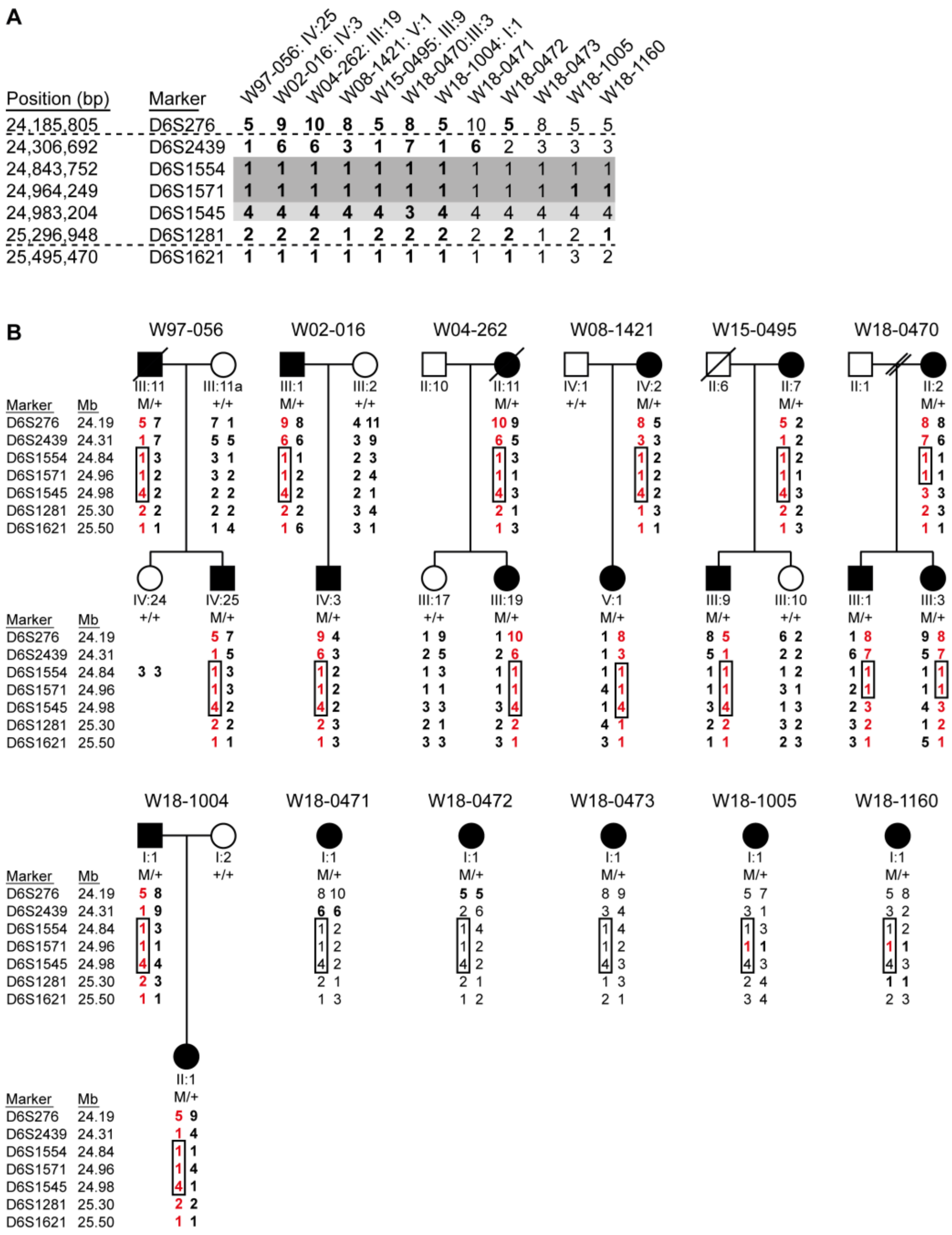


**Supplemental Figure 4. Family pedigrees with genotypes and haplotypes of VNTR markers.** (A) The shared haplotype is marked in grey. For marker D6S1545 (light-grey), a different CA-repeat length was determined in one family, but the marker is considered to be potentially part of the shared haplotype as a change of repeat length cannot be excluded. Markers for which the phase of the alleles could conclusively be determined via segregation in the family are marked in bold. The *RIPOR2* c.1696_1707del variant is located between the markers D6S2439 and D6S1554. Genomic positions (bp) are according to the UCSC Genome Browser (GRCh37/hg19). (B) The haplotypes carrying the *RIPOR2* c.1696_1707del variant are shown in red. A haplotype of 1.1 Mb was found to be shared (boxed) and is delimited by the markers D5S1554 and D6S1545. Alleles for which the parent of origin could conclusively be determined are marked in bold.

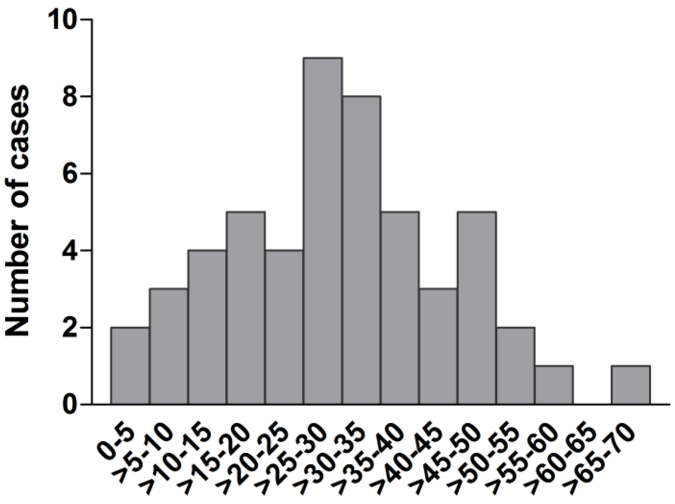


**Supplemental Figure 5. Reported age of onset of hearing loss of subjects with the *RIPOR2* c.1696_1707del variant.** Distribution of the reported ages of onset of *RIPOR2*-associated hearing loss per 5 years for 52 subjects who reported a specific age of onset. y, years.

# W97-056

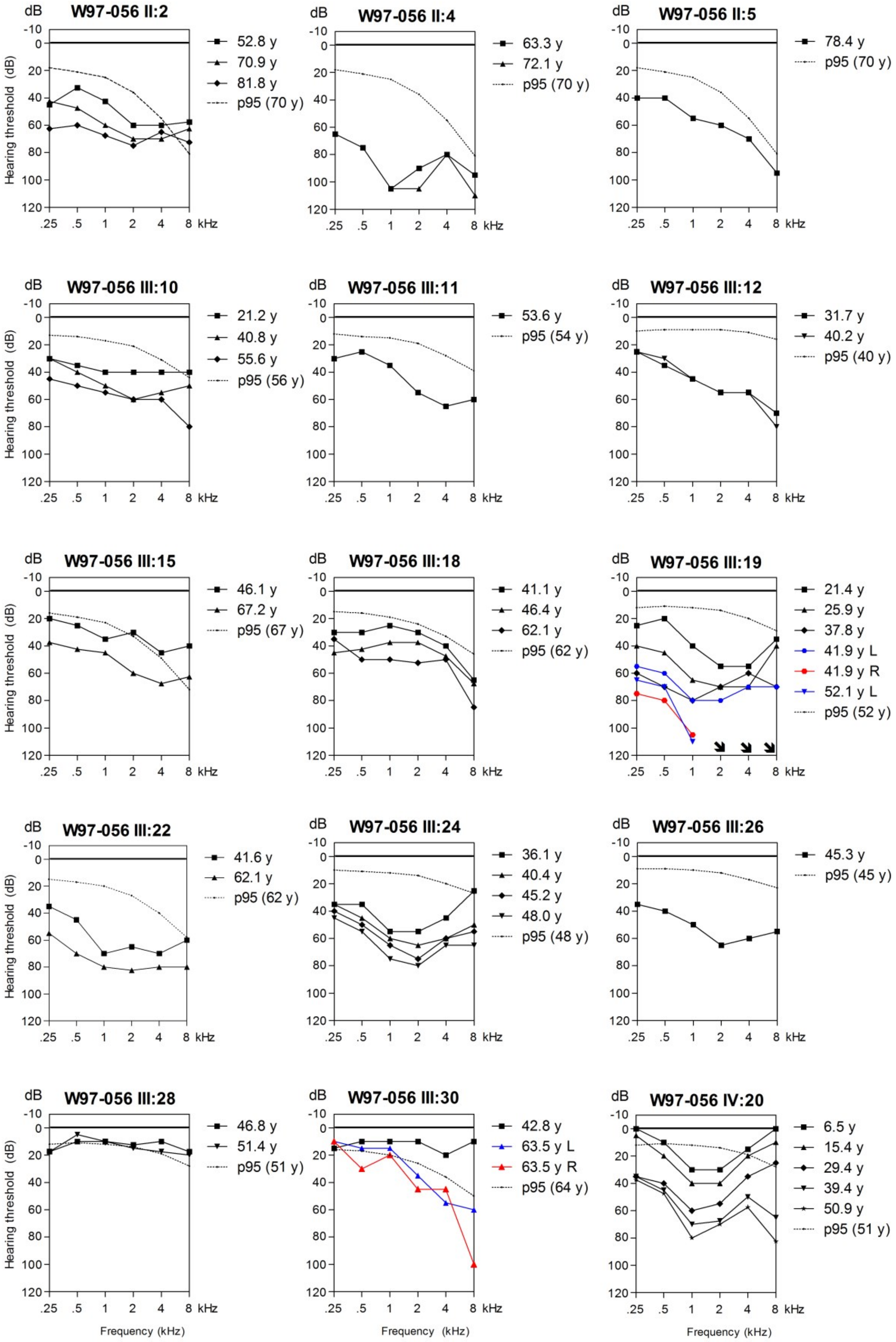

W97-056 II:2
52.8 y
70.9 y
81.8 y
p95 (70 y)
W97-056 II:4
63.3 y
72.1 y
p95 (70 y)
W97-056 II:5
78.4 y
p95 (70 y)
W97-056 III:10
21.2 y
40.8 y
55.6 y
p95 (56 y)
W97-056 III:11
53.6 y
p95 (54 y)
W97-056 III:12
31.7 y
40.2 y
p95 (40 y)
W97-056 III:15
46.1 y
67.2 y
p95 (67 y)
W97-056 III:18
41.1 y
46.4 y
62.1 y
p95 (62 y)
W97-056 III:19
21.4 y
25.9 y
37.8 y
41.9 y L
41.9 y R
52.1 y L
p95 (52 y)
W97-056 III:22
41.6 y
62.1 y
p95 (62 y)
W97-056 III:24
36.1 y
40.4 y
45.2 y
48.0 y
p95 (48 y)
W97-056 III:26
45.3 y
p95 (45 y)
W97-056 III:28
46.8 y
51.4 y
p95 (51 y)
W97-056 III:30
42.8 y
63.5 y L
63.5 y R
p95 (64 y)
W97-056 IV:20
6.5 y
15.4 y
29.4 y
39.4 y
50.9 y
p95 (51 y)
dB
Hearing threshold (dB)
-10
0
20
40
60
80
100
120
.25
.5
1
2
4
8
kHz
Frequency (kHz)

# W97-056

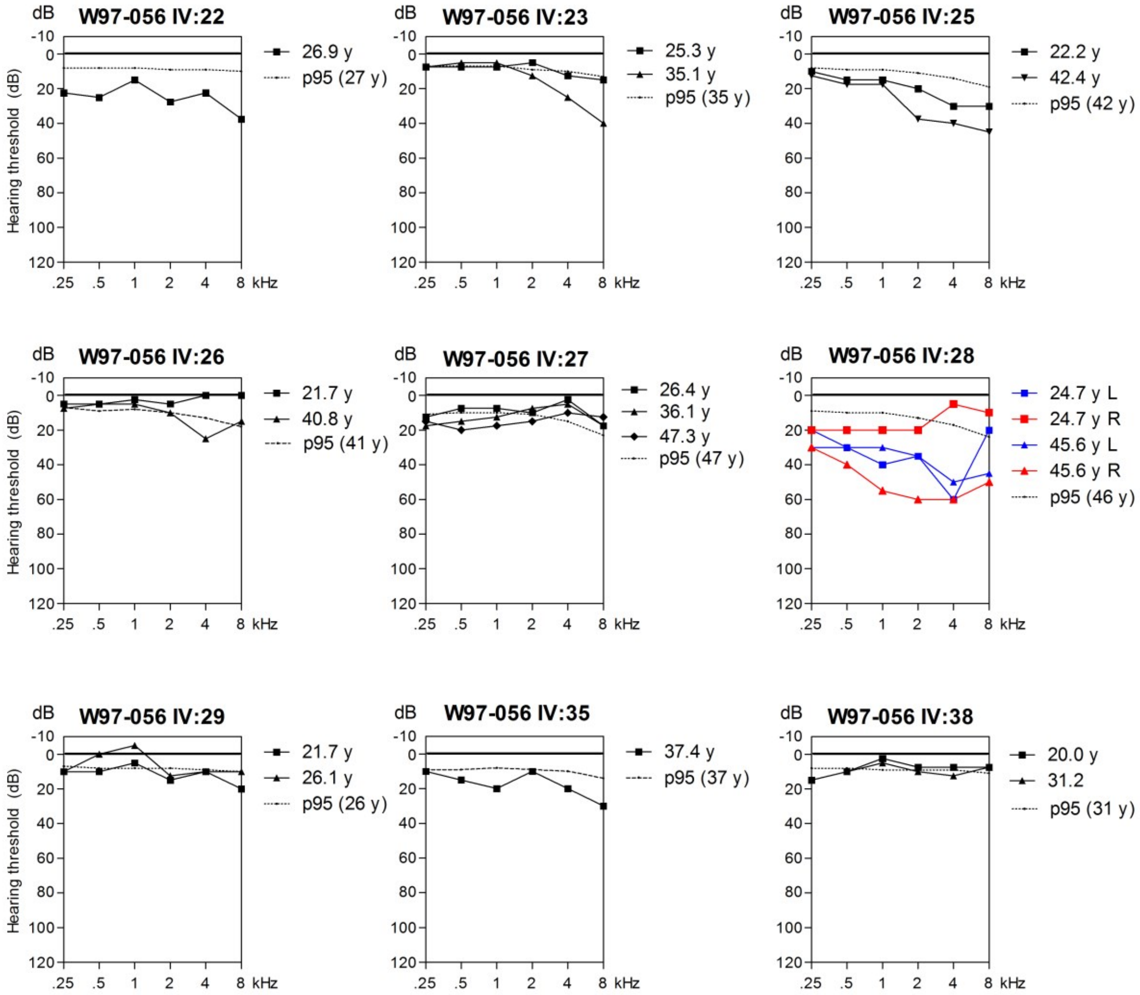


# W97-056, W02-016

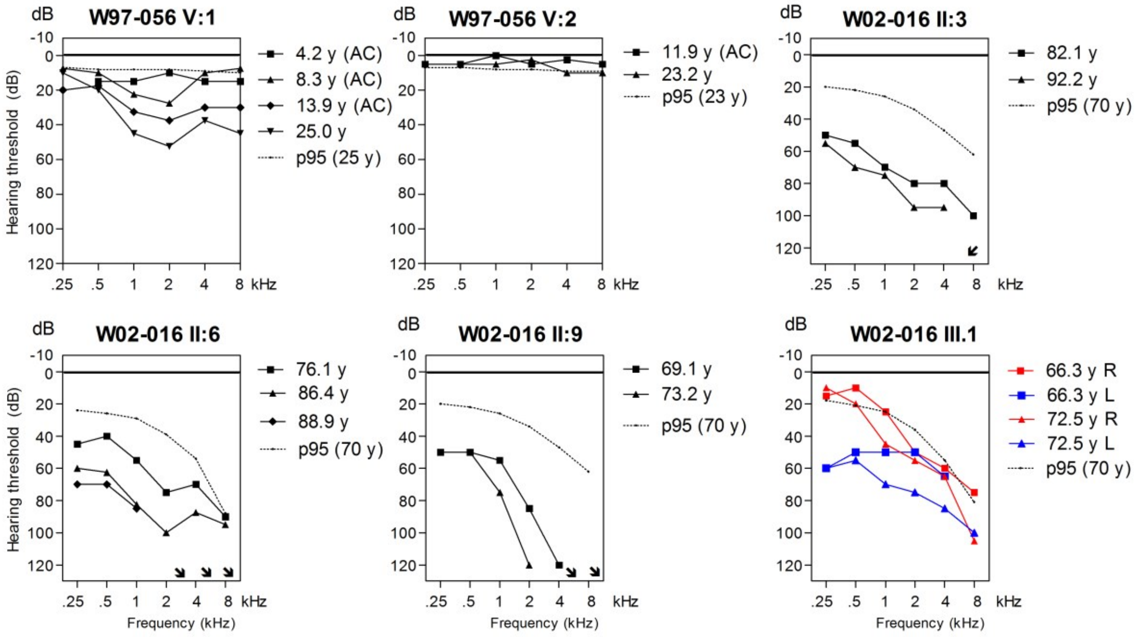

## W02-016

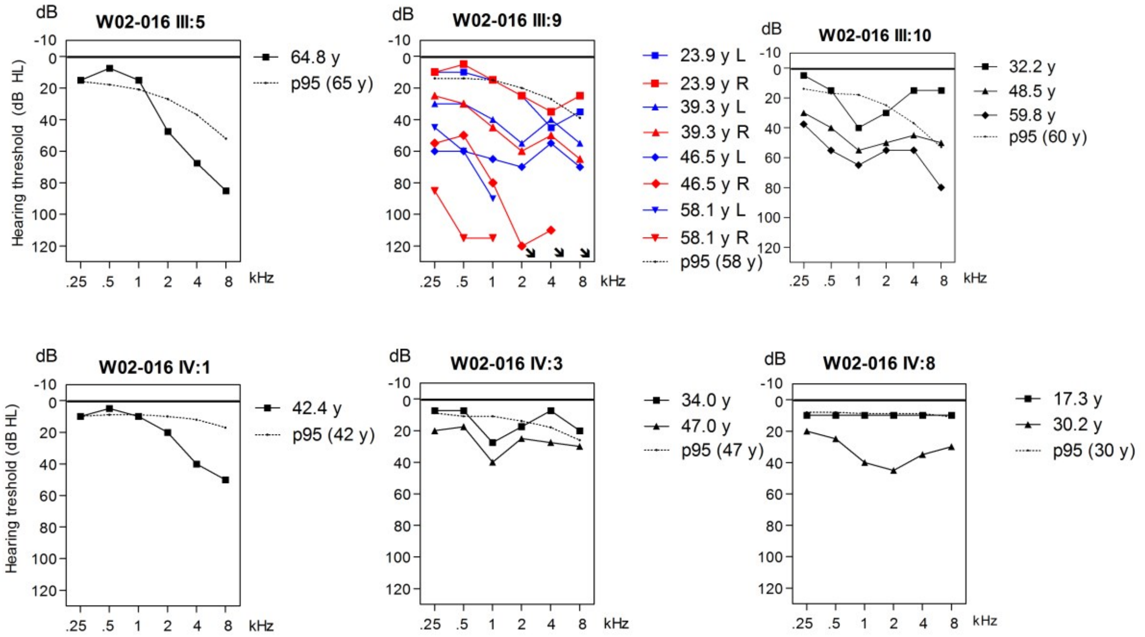


## W04-262

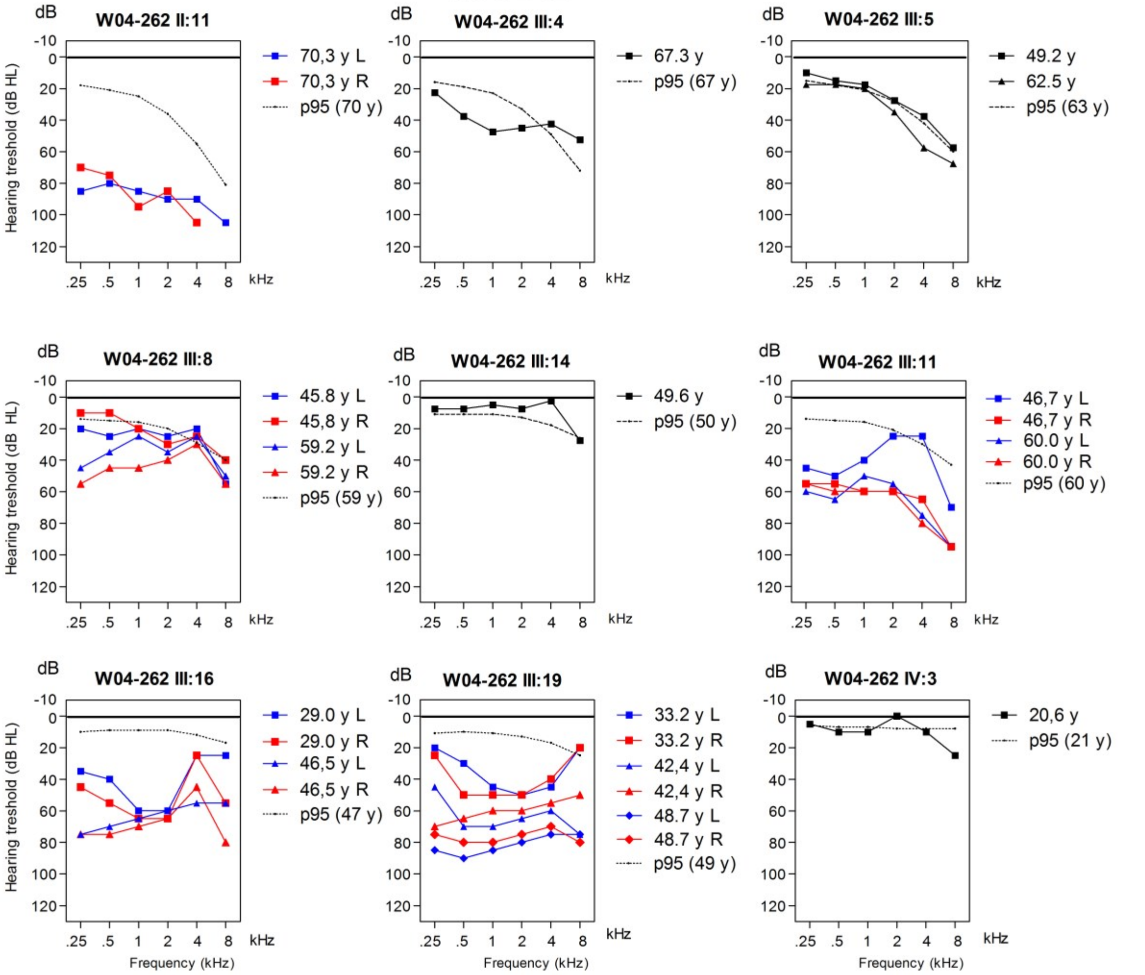

# W15-0495

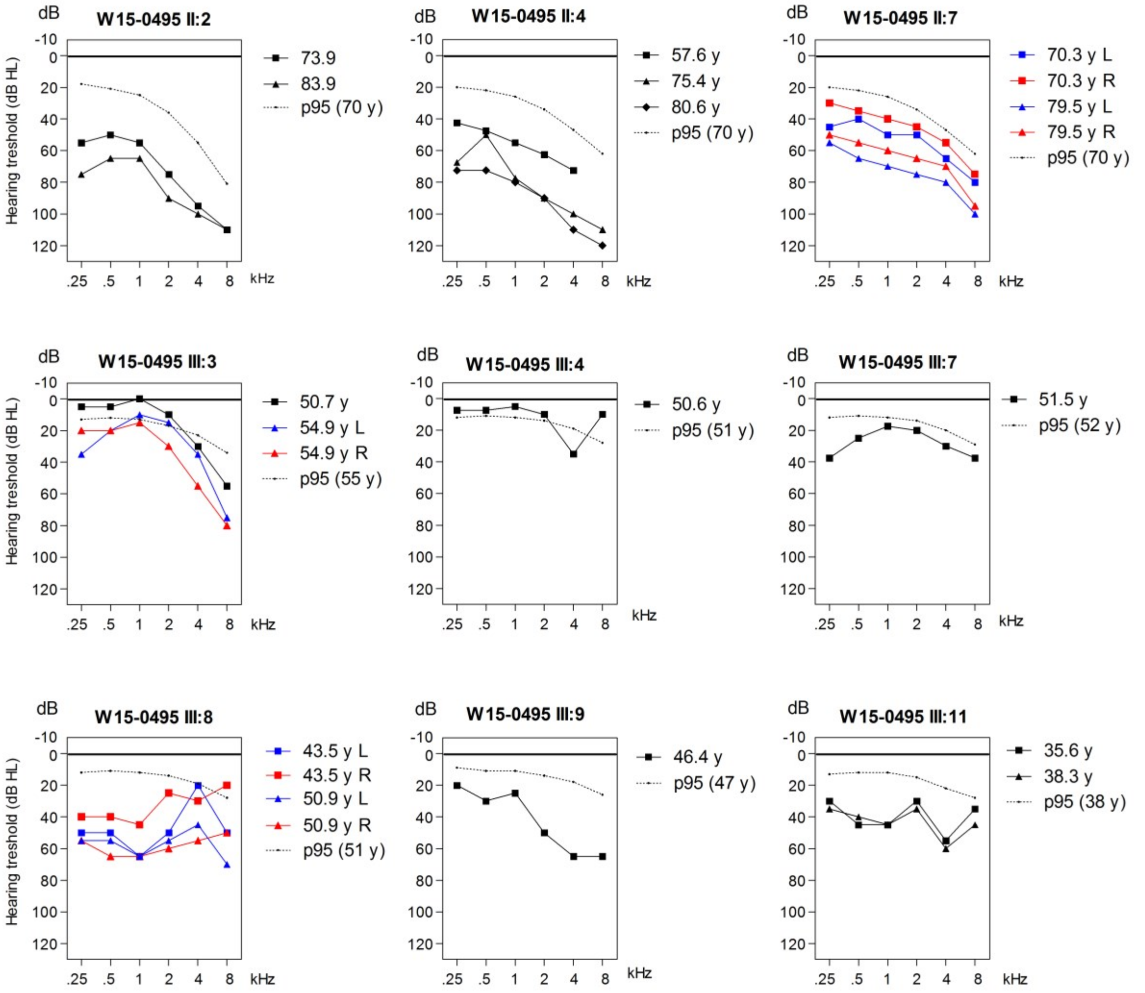


# W08-1421

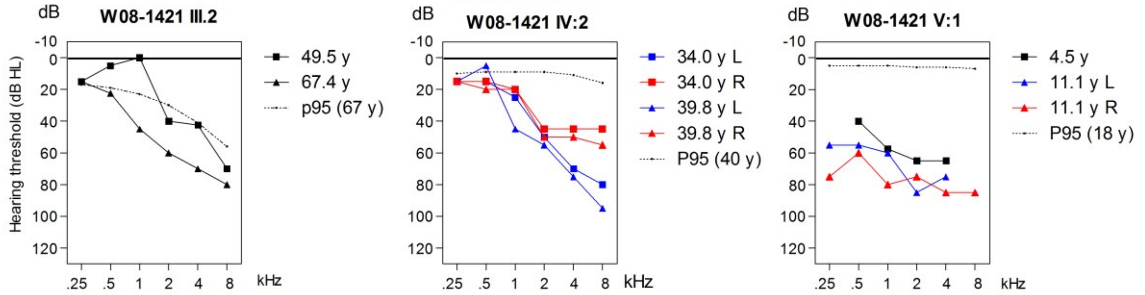


# W18-0470

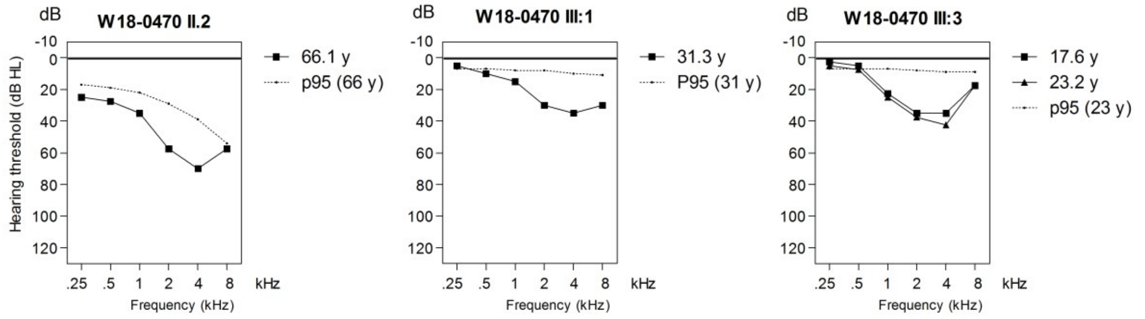

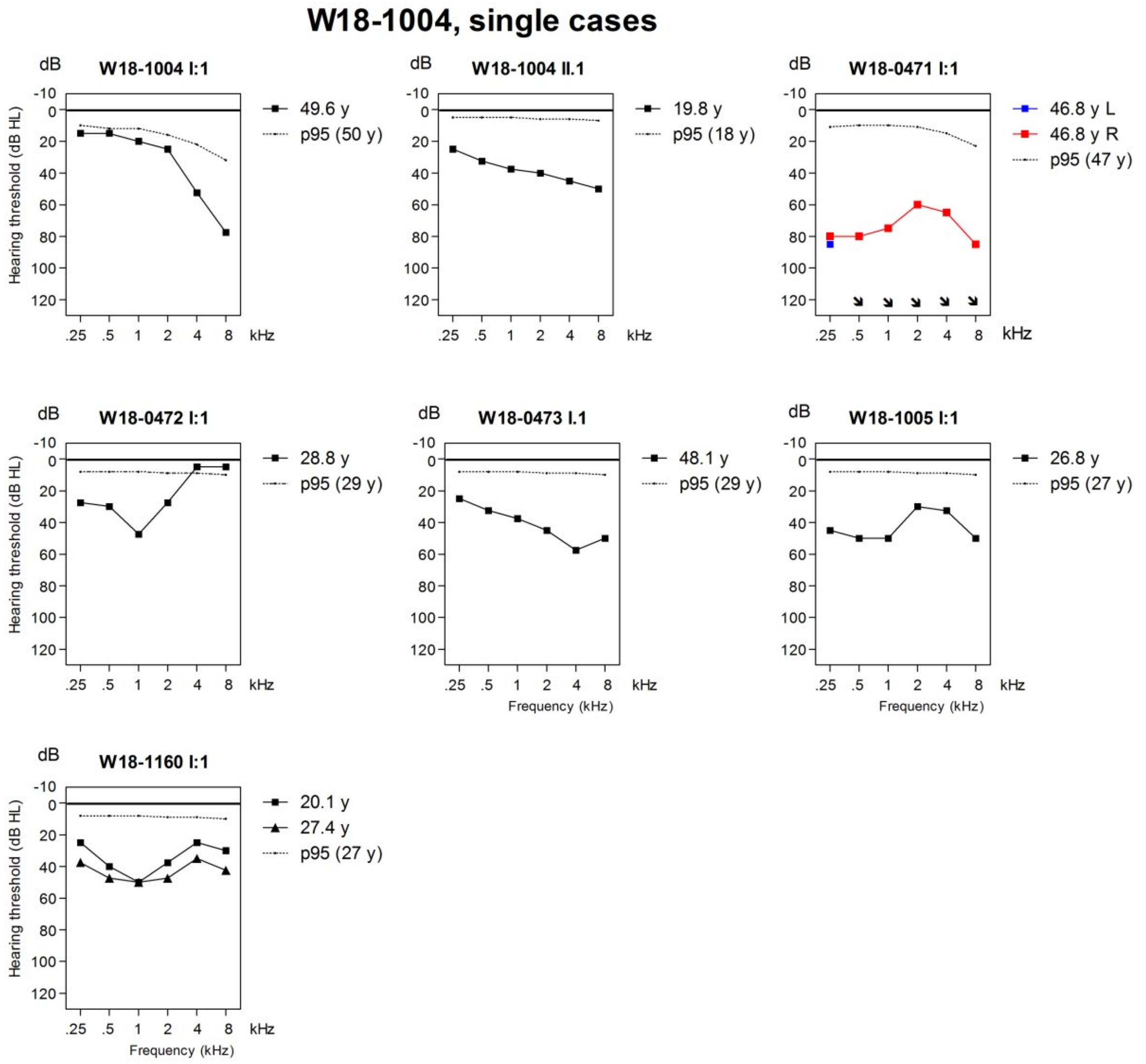


**Supplemental Figure 6. Pure tone audiometry.** Air conduction thresholds of all subjects (except III.5 of W08-1421) with the c.1696_1707del *RIPOR2* variant are depicted. In case of symmetry, the averages of left and right ear thresholds are shown. Otherwise, colorized (right red, left blue) audiograms of both ears are depicted. The p95 values are matched to the individuals' sex and age at the most recent audiometry, according to the ISO 7029:2017 standard. The age range for which the ISO 7029:2017 can be applied is 18 to 70 years. Part of the pure tone audiometry for family W97-056 has been published previously (9). y, age in years; R, right; L, left; dB HL, decibel hearing level; kHz, kiloHertz; AC; only air conduction levels available, no additional bone conduction thresholds have been measured.

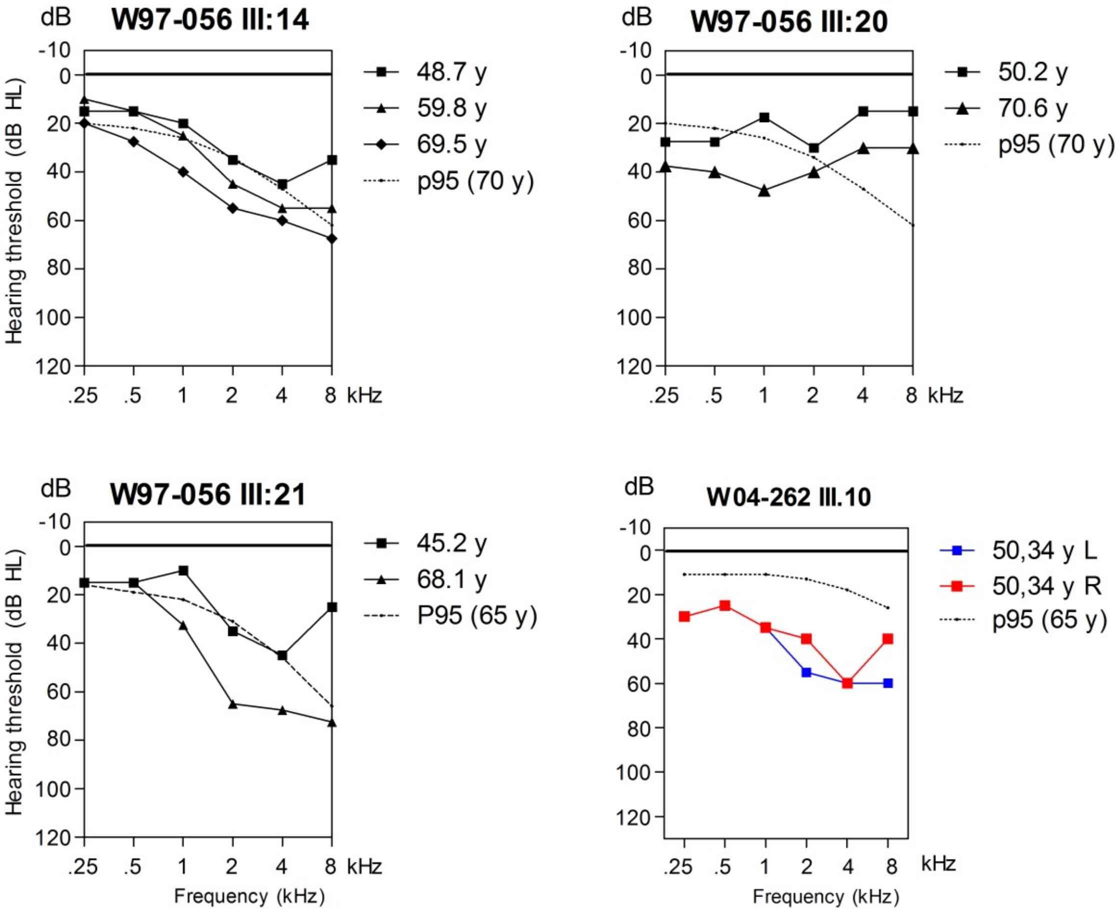


**Supplemental Figure 7. Audiograms of hearing impaired individuals without the *RIPOR2* c.1696_1707del variant.** Air conduction thresholds are depicted of all hearing impaired subjects who did not have the c1696_1707del *RIPOR2* variant. In supplemental table 5 information about possible explanations for their hearing impairment is provided. In case of symmetry, the averages of left and right ear thresholds are shown. Otherwise, colorized (right red, left blue) audiograms of both ears are depicted. The p95 values are matched to the individuals' sex and age at the most recent audiometry, according to the ISO 7029:2017 standard. The age range for which the ISO 7029:2017 can be applied is 18 to 70 years. y, age in years; R, right; L, left; dB HL, decibel hearing level; kHz, kiloHertz.

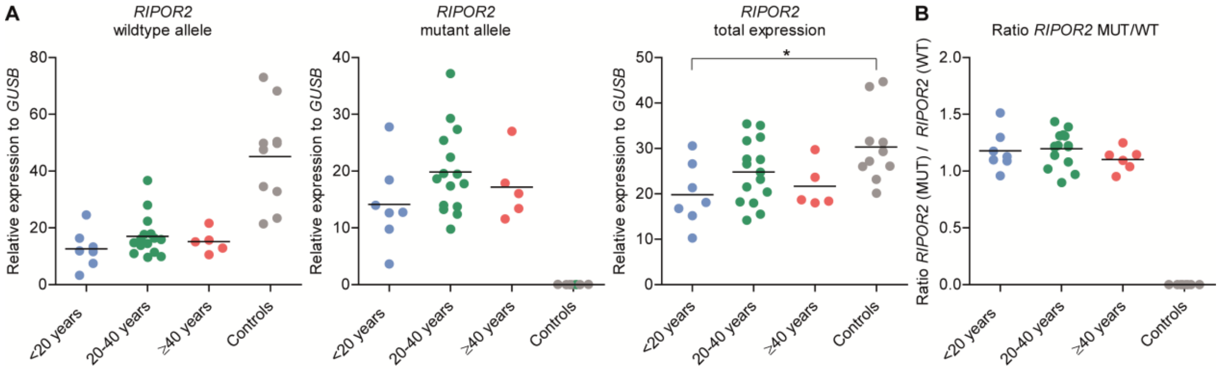


**Supplemental Figure 8. Transcript levels of *RIPOR2* alleles determined by RT-qPCR.** (A) Subjects were divided in three groups based on the reported ages of onset: early onset (<20 years, n=7), middle onset, (20-39 years, n=15) and late onset (≥40, n=6) hearing impairment. RNA samples isolated from peripheral blood of individuals without the *RIPOR2* variant were used as controls (n=10). (B) Calculated ratio of *RIPOR2* mutant to wildtype relative expression analysis. A one-way ANOVA followed by Tukey's multiple comparison test was employed to identify potentially significant differences between the transcript levels of the groups. ***** p-value = 0.0214.

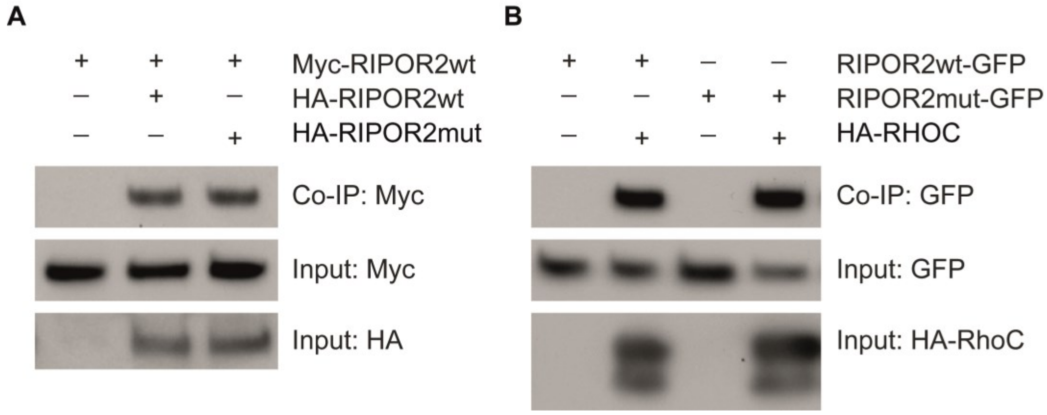


**Supplemental Figure 9. RIPOR2 imerization and interaction with RHOC.** (A) Interaction of murine RIPOR2-wildtype (RIPOR2wt) and -mutant (RIPOR2mut) was studied using CoIP assays. HEK293T cells were transfected with constructs encoding N-terminally tagged proteins as indicated above each panel. Immunoprecipitations were performed using anti-HA antibodies, followed by western blotting. (B) Interaction with RHOC was studied using C-terminally GFP-tagged RIPOR2-wildtype or -mutant and N-terminally HA-tagged RHOC. Immunoprecipitations were performed using anti-HA antibodies, followed by western blotting.

**Supplemental Table 1. Shared rare WES variants in family W97-056**

| Genome | Gene | Transcript | cDNA | Protein | In-house AF (%) | gnomAD_E AF (%) | gnomAD_G AF (%) | CADD_ PHRED | SIFT | PPH2 | MutationTaster (prob) |
|---|---|---|---|---|---|---|---|---|---|---|---|
| Chr1: 248,059,798T>A | *OR2W3* | NM_001001957.2 | c.910T>A | p.(Leu304Met) | 0.07 | 0.02 | 0.03 | 6.504 | 0.1 | 0.032 | Polymorphism (1.0) |
| Chr6: 15,501,310C>T | *JARID2* | NM_004973.3 | c.2118C>T | p.(Leu706=) | 0.18 | 0.03 | 0.04 | NA | NA | NA | NA |
| Chr6: 16,146,884C>T | *MYLIP* | NM_013262.3 | c.1249-9C>T | - | 0.01 | - | - | NA | NA | NA | NA |
| Chr6: 24,843,303_24,843,314del | *RIPOR2* | NM_014722.3 | c.1696_1707del | p.(Gln566_Lys569del) | 0.08 | 0.00 | - | NA | NA | NA | NA |
| Chr6: 41,196,733C>T | *TREML4* | NM_198153.2 | c.345C>T | p.(Ser115=) | 0.47 | 0.31 | 0.30 | NA | NA | NA | NA |
| Chr6: 44,329,574G>A | *SPATS1* | NM_145026.3 | c.419G>A | p.(Gly140Glu) | 0.18 | 0.11 | 0.12 | 19.16 | 0.02 | 1.0 | Disease causing (0.98) |

Variants identified by whole exome sequencing (WES) that are shared by all three index cases of family W97-057 and have an allele frequency of ≤0.5% in gnomAD and the in-house database (~7,500 exomes). For none of the variants is an effect on transcript splicing predicted nor are any reported in the ClinVar database. Scores that meet the thresholds for pathogenicity as described in the methods section are indicated in red. Thresholds for pathogenicity: CADD-PHRED (≥15), SIFT (≤0.05), PolyPhen-2 (≥0.450) and MutationTaster (disease causing). Genome, Genomic positions according to GRCh37/hg19; In-house AF, allele frequency (%) in the in-house database (~15,000 alleles); GnomAD_E AF and GnomAD_G AF, allele frequencies (%) in respectively gnomAD exome or genome databases; CADD_PHRED, Combined Annotation Dependent Depletion PHRED score; SIFT, Scale-Invariant Feature Transform; PPH2, Poly-Phen-2 score; MutationTaster (prob), MutationTaster score with probability (0-1); -, frequency not available; NA, not applicable.

**Supplemental Table 2. Rare variants in the index cases in genes known to be associated with adHL**

| Family | Genome | Gene | Transcript | cDNA | Protein | In-house AF (%) | gnomAD_E AF (%) | gnomAD_G AF (%) | CADD_ PHRED | SIFT | PPH2 | MutationTaster (prob) | ClinVar |
|---|---|---|---|---|---|---|---|---|---|---|---|---|---|
| W18-0470 | Chr22: 36681327T>C | *MYH9* | NM_002473.4 | c.5323A>G | p.(Lys1775Glu) | 0.18 | 0.15 | 0.19 | 22.2 | 0.03 | 0.120 | Disease causing (1.0) | UV2 |
| W18-0473 | Chr4: 6303119C>T | *WFS1* | NM_006005.3 | c.1597C>T | p.(Pro533Ser) | 0.22 | 0.07 | 0.08 | 19.64 | 0.00 | 1.000 | Disease causing (1.0) | UV1-UV3 |
| | Chr22: 36700183G>A | *MYH9* | NM_002473.4 | c.2248G>A | p.(Asp750Asn) | 0.01 | 0.00 | - | 20.80 | 0.00 | 0.997 | Disease causing (1.0) | NA |
| W18-1160 | Chr11: 76873225A>G | *MYO7A* | NM_000260.3 | c.1403A>G | p.(His468Arg) | 0.10 | 0.01 | 0.02 | 19.61 | 0.01 | 0.993 | Disease causing (1.0) | UV3 |

For none of the variants is an effect on transcript splicing predicted. Scores that meet the thresholds for pathogenicity as described in the methods section are indicated in red. Thresholds for pathogenicity: CADD-PHRED (≥15), SIFT (≤0.05), PolyPhen-2 (≥0.450) and MutationTaster (deleterious). Genome, Genomic positions according to GRCh37/hg19; In-house AF, allele frequency (%) in in-house database (~7,500 exomes); GnomAD_E AF and GnomAD_G AF, allele frequencies (%) in respectively gnomAD exome or genome databases; CADD_PHRED, Combined Annotation Dependent Depletion PHRED score; SIFT, Scale-Invariant Feature Transform; PPH2, Poly-Phen-2 score; MutationTaster (prob), MutationTaster score with probability (0-1); ClinVar, American College of Medical Genetics and Genomics (ACMG) classification of variants as in ClinVar; UV1, benign; UV2, likely benign; UV3, variant with unknown significance; NA, not available.

**Supplemental Table 3. Individual results of otoscopic examination, audiometry, imaging and progression of HL**

| Family | Subject | Age of onset (y) | Otoscopic examination | Clinical remarks | Imaging | | Audiometry | | | | | | | Progression of HL | | General remarks |
|---|---|---|---|---|---|---|---|---|---|---|---|---|---|---|---|---|
| | | | | | | | Subject age (y) | PTA | | SRT | | Maximum SRS (%) | | Progression rate (dB/y) | YOF (y) | |
| | | | | | CT | MRI | | R | L | R | L | R | L | | | |
| W97-056 | II:2 | 48 | NT | | | | 81 | 72 | 63 | 70 | 72 | 87 | 65 | 0.6 | 53-82 | |
| | II:4 | 43 | NT | Ab, NE, T | | | 72 | 93 | 90 | NT | NT | NT | NT | NA | 63-72 | |
| | II:5 | 70 | NT | NE | | | 79 | 63 | 52 | NT | NT | NT | NT | NA | 0 | Professional noise exposure |
| | III:10 | 21 | NT | NE | | | 41 | 47 | 48 | 61 | 70 | 95 | 95 | 0.5 | 21-56 | |
| | III.11 | 26 | NT | O, T | | | 54 | 43 | 38 | NT | NT | NT | NT | NA | 0 | |
| | III:12 | 20 | NT | T | | | 41 | 40 | 43 | 43 | 42 | 90 | 95 | NA | 32-40 | |
| | III:15 | 37 | N | NE,T | | | 67 | 50 | 48 | 57 | 51 | 92 | 75 | 1.0 | 46-67 | |
| | III:18 | 36 | NT | T | | | 62 | 50 | 51 | 50 | NT | 95 | 90 | 0.9 | 41-62 | |
| | III:19 | 14 | N | A | | | 42 | NA | 73 | NA | NA | NA | 41 | 1.4 | 21-42 | R ear profoundly deaf |
| | III:22 | 20 | NT | T | | | 62 | 75 | 80 | 70 | 70 | 75 | 88 | 0.8 | 42-62 | |
| | III:24 | 33 | N | | | | 43 | 62 | 60 | 63 | 67 | 93 | 93 | 1.8 | 36-48 | |
| | III:26 | 34 | N | O,T | | | 45 | 50 | 52 | NT | NT | NT | NT | NA | 0 | |
| | III:28 | NOHL | NT | | | | 52 | 7 | 10 | NT | NT | NT | NT | NA | 47-51 | |
| | III:30 | NR | N | A | | | 64 | 53 | 35 | 42 | 25 | 100 | 100 | 1.2 | 48-64 | otosclerosis, infrequent balance complaints |
| | IV:20 | 7 | N | T | | | 50 | 70 | 63 | 62 | 52 | 88 | 90 | 0.9 | 7-51 | |
| | IV:22 | NR | N | T | | | 27 | 20 | 25 | NT | NT | NT | NT | NA | 0 | |
| | IV:23 | NR | NT | | | | 36 | 7 | 8 | NT | NT | NT | NT | 0.5 | 25-35 | |
| | IV:25 | 24 | NT | | | | 42 | 25 | 18 | 18 | 22 | NT | NT | 0.4 | 22-42 | |

| (Continued) Family | Subject | Age of onset (y) | Otoscopic examination | Clinical remarks | Imaging | | Audiometry | | | | | | | Progression of HL | | General remarks |
|---|---|---|---|---|---|---|---|---|---|---|---|---|---|---|---|---|
| | | | | | | | Subject age (y) | PTA | | SRT | | Maximum SRS (%) | | Progression rate (dB/y) | YOF (y) | |
| | | | | | CT | MRI | | R | L | R | L | R | L | | | |
| **W97-056** | IV:26 | NOHL | NT | | | | 41 | 7 | 7 | NT | NT | NT | NT | NA | NA | |
| | IV:27 | 35 | N | | | | 47 | 17 | 18 | 12 | 18 | 100 | 100 | 0.3 | 26-47 | |
| | IV:28 | 30 | N | A, NE, T | N | N | 45 | 32 | 52 | 37 | 55 | 100 | 92 | 1.8 | 25-46 | Professional noise exposure |
| | IV:29 | 17 | N | NE | | | 26 | 5 | 10 | NT | NT | NT | NT | NA | 22-26 | |
| | IV:35 | 30 | N | NE, T | | | 37 | 15 | 15 | 10 | 10 | 100 | 100 | NA | 0 | Recreational noise exposure |
| | IV:38 | 8 | N | T | | | 31 | 8 | 7 | 8 | 7 | 100 | 100 | NA | NA | |
| | V:1 | 5 | N | NE, T | | | 25 | 40 | 37 | 28 | 27 | 100 | 97 | 1.3 | 8-25 | |
| | V:2 | NOHL | N | | | | 23 | 3 | 5 | <10 | <10 | 100 | 100 | NA | NA | |
| **W02-016** | II:3 | 29 | NT | T | | | 82 | 68 | 68 | 65 | 70 | 100 | 93 | 0.9 | 82-92 | |
| | II:6 | 47 | N | | | | 89 | 80 | 83 | 80 | 80 | 52 | 67 | 2 | 76-86 | |
| | II:9 | 40 | N | | | | 73 | NA | 82 | 67 | 72 | 75 | 55 | NA | 69-73 | R ear profoundly deaf above 2 kHz |
| | III:1 | 41 | N | A, V | | | 73 | 40 | 67 | 35 | 77 | 88 | 62 | NA | 66-73 | Infrequent vertigo attacks since the age of 65 years |
| | III:5 | NR | N | | | | 65 | 20 | 27 | 18 | 17 | 95 | 96 | NA | 0 | |
| | III:9 | 35 | N | O, A, T, V | N | | 39 | 48 | 42 | 47 | 38 | 80 | 100 | 3 | 24-47 | Balance complaints after CI surgery |
| | III:10 | 32 | N | T | | | 32 | 15 | 20 | 10 | 15 | 100 | 95 | 1.4 | 32-60 | |
| | IV:1 | 53 | NT | V | | | 42 | 12 | 10 | NT | NT | NT | NT | NA | 0 | Benign paroxymal positional vertigo |
| | IV:3 | 17 | N | | | | 47 | 27 | 25 | 25 | 17 | 100 | 100 | 1 | 34-47 | |
| | IV:8 | 27 | N | | | | 30 | 38 | 37 | 32 | 30 | 100 | 95 | 2.7 | 17-30 | |

| (Continued) Family | Subject | Age of onset (y) | Otoscopic examination | Clinical remarks | Imaging | | Audiometry | | | | | | | Progression of HL | | General remarks |
|---|---|---|---|---|---|---|---|---|---|---|---|---|---|---|---|---|
| | | | | | | | Subject age (y) | PTA | | SRT | | Maximum SRS (%) | | Progression rate (dB/y) | YOF (y) | |
| | | | | | CT | MRI | | R | L | R | L | R | L | | | |
| **W04-262** | II:11 | <18 | R atelectasis L sclerotic | A, T | | | 70 | 85 | 87 | >95 | >95 | 37 | 44 | NA | 0 | Multiple ear surgeries, a.o. ear drum surgery |
| | III:4 | 49 | N | T | | | 67 | 40 | 47 | 28 | 35 | 100 | 90 | NA | 0 | |
| | III:5 | 55 | N | NE, T | | | 63 | 32 | 23 | NT | NT | NT | NT | 0.6 | 49-63 | |
| | III:8 | 49 | N | A | | | 59 | 43 | 32 | 37 | 22 | 100 | 100 | 1.4 | 46-59 | |
| | III:11 | 32 | N | A, T | | N | 60 | 62 | 55 | 60 | 70 | 95 | 70 | 2 | 47-60 | |
| | III:14 | NOHL | N | Ab | | | 49 | 7 | 7 | NT | NT | NT | NT | NA | NA | |
| | III:16 | 25 | N | A, T | | | 47 | 70 | 65 | 80 | 77 | 55 | 60 | 0.9 | 29-47 | |
| | III:19 | 28 | N | A | | N | 49 | 83 | 92 | NA | NA | 42 | 42 | 2.1 | 33-49 | |
| | IV:3 | NR | N | | | | 21 | 12 | 7 | NT | NT | NT | NT | NA | 0 | |
| **W15-0495** | II:2 | 35 | NT | T | | | 84 | 73 | 75 | 65 | 70 | 60 | 70 | 0.9 | 74-84 | |
| | II:4 | PS | N | | | | 81 | 77 | 85 | 105 | 105 | 60 | 50 | 2.4 | 58-81 | |
| | II:7 | 60 | NT | A | | | 80 | 60 | 73 | 55 | 72 | 73 | 55 | 1.5 | 70-80 | |
| | III:3 | 38 | N | T, A | | | 55 | 22 | 15 | 19 | 14 | 100 | 97 | NA | 51-55 | |
| | III:4 | NOHL | NT | NE | | | 51 | 8 | 7 | NT | NT | NT | NT | NA | 0 | Noise trauma |
| | III:7 | 50 | NT | | | | 52 | 22 | 20 | NT | NT | NT | NT | NA | 0 | |
| | III:8 | 30 | N | A, T | | | 48 | 62 | 58 | 52 | 48 | 92 | 92 | NA | 44-51 | |
| | III:9 | 43 | N | NE | | | 46 | 38 | 37 | 42 | 44 | 96 | 90 | NA | 0 | Professional noise exposure |
| | III:11 | 33 | N | NE, T | | | 38 | 38 | 37 | 40 | 40 | 100 | 95 | NA | 36-38 | |

| (Continued) Family | Subject | Age of onset (y) | Otoscopic examination | Clinical remarks | Imaging | | Audiometry | | | | | | | Progression of HL | | General remarks |
|---|---|---|---|---|---|---|---|---|---|---|---|---|---|---|---|---|
| | | | | | | | Subject age (y) | PTA | | SRT | | Maximum SRS (%) | | Progression rate (dB/y) | YOF (y) | |
| | | | | | CT | MRI | | R | L | R | L | R | L | | | |
| **W08-1421** | III:2 | cong. | N | T | | | 67 | 42 | 43 | 30 | 40 | 87 | 65 | 1.8 | 50-67 | |
| | IV:2 | PS | N | A, Ns, O, T | | | 34 | 27 | 28 | 27 | 32 | 95 | 93 | NA | 6 | |
| | V:1 | 0 | R eardrum perforation | A, Ab, Ns, O, T | | | 9 | 73 | 67 | 58 | 60 | NT | NT | NA | NA | Neonatal intensive care, surgeries, ototoxic antibiotics |
| **W18-0470** | II:2 | 6 | N | | | | 66 | 42 | 38 | 27 | 27 | 100 | 95 | NA | 0 | |
| | III:1 | 30 | N | O | | | 31 | 19 | 18 | 16 | 14 | 100 | 100 | NA | 0 | |
| | III:3 | 11 | NT | | | | 23 | 23 | 22 | 15 | 13 | 100 | 100 | NA | 5 | |
| **W18-1004** | I:1 | 12 | N | | | | 54 | 17 | 23 | NT | NT | NT | NT | NA | 0 | |
| | II:1 | 4 | N | | | | 20 | 38 | 35 | 30 | 27 | 100 | 95 | NA | 5 | |
| **W18-0471** | I:1 | 20 | N | A, T | | | 47 | 72 | 117 | 64 | NA | 83 | 0 | NA | 0 | |
| **W18-0472** | I:1 | 27 | N | T, V | | | 31 | 35 | 37 | 18 | 22 | 100 | 100 | NA | 0 | Migrainous vertigo |
| **W18-0473** | I:1 | 39 | NT | | N | N | 48 | 35 | 42 | NT | NT | NT | NT | NA | 0 | |
| **W18-1005** | I:1 | 25 | N | | | N | 26 | 42 | 45 | 32 | 37 | 100 | 97 | NA | 0 | |
| **W18-1160** | I:1 | 12 | N | T | | | 50 | 47 | 48 | 42 | 43 | 100 | 100 | NA | 7 | |

Age of onset (AoO), age of onset in years as reported by the subjects. Subject age, the age at which the audiometric data of column 9 to 14 were obtained, in general the last audiogram. If no speech audiometry was performed during the latest pure tone audiometry, the latest audiogram in which both were measured, was selected. Progression rate of HL, calculated as described in the methods section, if there was at least a follow-up duration of 10 years. Y, years; PTA, pure tone average, mean of 0.5, 1 and 2 kHz air conduction thresholds; R, right; L, left; SRT, speech reception threshold; SRS, speech recognition score in %; YOF, years of follow up; NOHL, unaffected subject; NR, age of onset of HL not reported; PS, subject reported onset of HL during primary school; NT, not tested; N, no abnormalities; T, Tinnitus; NE, extensive exposure to noise; O, recurrent otitis; A, asymmetric HL; Ab, subject reported long-term antibiotics usage, but no details about duration and which antibiotics; V, vestibular complaints; NA, not applicable.

**Supplemental Table 4. Results of vestibular testing**

| Family | Subject (age) | Click-evoked ABR | Remarks and history of vestibular symptoms | Oculu-motor testing | vHIT (gain) | SPV Caloric irrigation: Warm (°/s) (10-52)[a] | | Cold (°/s) (7-31)[a] | | Conclusion | Rotating chair: Gain (%) (33-72)[a] | | SPV (°/s) (30-65)[a] | | Tau (s) (11-26)[a] | | Conclusion |
|---|---|---|---|---|---|---|---|---|---|---|---|---|---|---|---|---|---|
| | | | | | | R | L | R | L | | CW | CCW | CW | CCW | CW | CCW | |
| W97-056 | III:24 (48) | NT | no | normal | NT | NT | NT | 12 | 13 | normal | NA | NA | 49 | 50 | 15 | 15 | normal |
| | III:11 (53) | NT | no | normal | NT | 31 | 18 | 16 | 9 | normal | NA | NA | 28 | 30 | 14 | 14 | Underestimated[b] |
| W02-016 | III:1 (71) | N | Infrequent vertigo attacks since the age of 65 years | normal | NT | NT | NT | 3 | 11 | hyporeactive | NA | NA | 51 | 33 | 10 | 11 | Hyporeactive |
| | III:9 (63)[c] | N | Balance complaints after CI surgery | normal | normal | NT | NT | 19 | 24 | normal | 70 | 61 | 63 | 56 | 15 | 19 | normal |
| | III:10 (60) | N | no | normal | normal | 32 | 39 | NT | NT | normal | 46 | 57 | 42 | 52 | 12 | 12 | normal |
| W04-262 | III:16 (60) | NT | no | normal | normal | 31 | 30 | 37 | 46 | normal | NA | NA | 42 | 65 | 22 | 15 | normal |
| | III:19 (48) | NT | no | normal | NT | 9 | 7 | 9 | 8 | normal | 65 | 78 | 59 | 71 | 21 | 15 | normal |
| W08-1421 | IV:2 (47) | NT | no | normal | normal | 23 | 19 | 12 | 19 | normal | 75 | 80 | 68 | 72 | 11 | 11 | normal |
| W18-0470 | III:1 (32) | N | no | normal | normal | 12 | 26 | 12 | 12 | normal | 82 | 60 | 75 | 55 | 10 | 12 | normal |

ABR, auditory brainstem response; vHIT, video head impulse test; °/s,degrees per seconds; SPV, slow phase velocity; Tau, time constant; R, right ear; L, left ear; CW, clock-wise; CCW, counter clock-wise; N, no abnormalities; NT, not tested; NA, not applicable. [a], normative values at our institute [b], nystagmus was suppressed due to stress [c], Subject was tested after CI surgery and also had c- and oVEMP testing, no abnormalities were objectified (data not shown).

**Supplemental Table 5. Individual results of age of onset, otoscopy, audiometry and progression of HL of the phenocopies**

| Family | Subject | Age of onset (y) | Otoscopic examination | Clinical remarks | Imaging | | Audiometry | | | | | | | Progression of HL | | General remarks |
|---|---|---|---|---|---|---|---|---|---|---|---|---|---|---|---|---|
| | | | | | | | Subject age (y) | PTA | | SRT | | Maximum SRS (%) | | Progression rate (dB/y) | YOF (y) | |
| | | | | | CT | MRI | | R | L | R | L | R | L | | | |
| W97-056 | III:14 | 52 | N | V | | | 69 | 42 | 40 | 48 | 42 | 85 | 92 | 0.8 | 39-70 | Ménière-like phenotype |
| | III:20 | 46 | N | Ab, T | | | 70 | 45 | 40 | 37 | 37 | 100 | 100 | 0.7 | 50-71 | Smoking, COPD Gold III, often antibiotics |
| | III:21 | NR | NT | | | | 68 | 38 | 37 | 20 | 25 | NT | NT | 0.8 | 45-68 | |
| W04-262 | III:10 | 55 | N | T, V | | | 60 | 33 | 38 | 33 | 41 | 100 | 95 | NA | 0 | |

Family members with HL that do not carry the *RIPOR2* variant are considered a phenocopy. Age of onset (AoO) is the age of onset in years as reported by the subjects. Subject age is the age at which the audiometric data of column 9 to 14 were obtained. If no speech audiometry was performed during the latest audiometric testing, the penultimate audiogram was selected. Progression rate of HL was calculated if there was at least a follow-up duration of 10 years. Y, years; PTA, pure tone average, mean of 0.5, 1 and 2 kHz air conduction thresholds; R, right; L, left; SRT, speech reception threshold; SRS, speech recognition score in %; YOF, years of follow up; NR, age of onset of HL not reported; NT, not tested; N, no abnormalities; V, vestibular complaints; Ab, subject reported long-term antibiotics usage, but no details about duration and which antibiotics; T, Tinnitus; NA, not applicable.

**Supplemental Table 6. Genes analyzed by MIP sequencing**

*ACTG1*

*ADCY1*

*BDP1*

*BSND*

*CABP2*

*CCDC50*

*CDH23*

*CEACAM16*

*CIB2*

*CLDN14*

*CLIC5*

*COCH*

*COL11A2*

*COL4A6*

*CRYM*

*DCDC2*

*GSDME*

*DFNB31*

*DFNB59*

*DIABLO*

*DIAPH1*

*ELMOD3*

*EPS8*

*ESPN*

*ESRRB*

*EYA4*

*GIPC3*

*GJB2*

*GJB3*

*GJB6*

*GPSM2*

*GRHL2*

*GRM7*

*GRM8*

*GRXCR1*

*GRXCR2*

*HGF*

*ILDR1*

*KARS*

*KCNQ4*

*LHFPL5*

*LOXHD1*

*LRTOMT*

*MARVELD2*

*MIR96*

*MSRB3*

*MYH14*

*MYH9*

*MYO15A*

*MYO3A*

*MYO6*

*MYO7A*

*NAT2*

*OSBPL2*

*OTOA*

*OTOF*

*OTOG*

*OTOGL*

*P2RX2*

*PCDH15*

*PNPT1*

*POU3F4*

*POU4F3*

*PRPS1*

*PTPRQ*

*RDX*

*RIPOR2*

*SERPINB6*

*SIX1*

*SLC17A8*

*SLC26A4*

*SLC26A5*

*SMPX*

*STRC*

*SYNE4*

*TBC1D24*

*TECTA*

*TJP2*

*TMC1*

*TMEM132E*

*TMIE*

*TMPRSS3*

*TNC*

*TPRN*

*TRIOBP*

*TSPEAR*

*USH1C*

*USH1G*

*WFS1*

**Supplemental Table 7. Primer sequences**

| Target | Primer | Oligonucleotides (5'-3') |
|---|---|---|
| *RIPOR2* exon 14, wt allele | Forward | aagcagctggtcaagagg |
| | Reverse | gcagccttcagattctcc |
| *RIPOR2* exon 14, mut allele | Forward | ggaaggaaacatcacaaagag |
| | Reverse | gcagccttcagattctcc |
| *RIPOR2* exons 3-4, mRNA | Forward | ggccttgaaaaatggacttg |
| | Reverse | ccaggcgagagtttcttttc |
| *RIPOR2*, exons 11-16, mRNA | Forward | accatcaaactgaacctgga |
| | Reverse | cccaactcctgtgtcttcag |
| *RIPOR2,* exons 12-15, mRNA | Forward | tccatgtacagccagggta |
| | Reverse | acttaaactggaagacctgctg |
| *GUSB* exons 2-3, mRNA | Forward | agagtggtgctgaggattgg |
| | Reverse | ccctcatgctctagcgtgtc |

Primer sequences for *RIPOR2* are based on reference sequence NM_00147722.3 and for *GUSB* on NM_00181.3. wt, wildtype; mut, mutant.

### SUPPLEMENTAL ACKNOWLEDGMENTS

The DOOFNL Consortium consists of the collaborators M.F. van Dooren, H.H.W. de Gier, E.H. Hoefsloot, M.P. van der Schroeff, S.G. Kant (ErasmusMC, Rotterdam, the Netherlands), L.J.C. Rotteveel, F.G. Ropers (LUMC, Leiden, the Netherlands), J.C.C. Widdershoven, J.R. Hof, E.K. Vanhoutte (MUMC+, Maastricht, the Netherlands), H. Kremer, R.J.E. Pennings, C.P. Lanting, H. Yntema, R.J.C. Admiraal, I. Feenstra (Radboudumc, Nijmegen, the Netherlands), R.H. Free and J.S. Klein Wassink-Ruiter (UMCG, Groningen, the Netherlands), R.J. Stokroos, A.L. Smit, M.J. van den Boogaard (UMC, Utrecht, the Netherlands) and F.A. Ebbens, S.M. Maas, A. Plomp, T.P.M. Goderie, P. Merkus and J. van de Kamp (Amsterdam UMC, Amsterdam, the Netherlands).
